# Natural variations in the P-type ATPase heavy metal transporter ZmCd1 controlling cadmium accumulation in maize grains

**DOI:** 10.1101/2021.01.13.426291

**Authors:** Bin Tang, Meijie Luo, Yunxia Zhang, Huanle Guo, Jingna Li, Wei Song, Ruyang Zhang, Zhen Feng, Mengsi Kong, Han Li, Zhongyang Cao, Xiaoduo Lu, Delin Li, Jianhua Zhang, Ronghuan Wang, Yuandong Wang, Zhihui Chen, Yanxin Zhao, Jiuran Zhao

## Abstract

Cadmium (Cd) accumulation in maize grains is detrimental to human health. Developing maize varieties with low-Cd contents via marker-assisted selection is important for ensuring the production of maize grains safe for consumption. However, the key gene controlling maize grain Cd accumulation has not been cloned. In this study, we identified two major loci for maize grain Cd accumulation (*qCd1* and *qCd2*) on chromosome 2 during a genome-wide association study (GWAS). The *qCd1* locus was analyzed by bulked segregant RNA-seq and fine mapping with a biparental segregating population of Jing724 (low-Cd line) and Mo17 (high-Cd line). The *ZmCd1* candidate gene in the *qCd1* locus encodes a vacuolar membrane-localized heavy metal P-type ATPase transporter, ZmHMA3, which is orthologous to the tonoplast Cd transporter OsHMA3. Genomic DNA sequence and transcript analyses suggested that a transposon in intron 1 of *ZmCd1* is responsible for the abnormal amino acid sequence in Mo17. An EMS mutant analysis and an allelism test confirmed *ZmCd1* influences maize grain Cd accumulation. The natural variations in *ZmCd1* were used to develop four PCR-based molecular markers, which revealed five *ZmCd1* haplotypes in the GWAS population. The molecular markers were also used to predict the grain Cd contents in commonly cultivated maize germplasms in China. The predicted Cd contents for 36 inbred lines and 13 hybrids were consistent with the measured Cd contents. Furthermore, several low-Cd elite inbred lines and hybrids were identified, including Jing2416, MC01, Jingnonke728, and Jingke968. Therefore, the molecular markers developed in this study are applicable for molecular breeding and developing maize varieties with low grain Cd contents.

## Introduction

Cadmium (Cd) is one of the most deleterious heavy metals. It can accumulate in the food chain, but even a very low Cd concentration can be fatal to humans because of its long half-life (Yan et al., 2019). Specifically, Cd can damage bones, the liver, kidneys, and lungs, and even cause cancer (Zhang et al., 2019). Unfortunately, Cd has been widely released into soils because of increasing industrial and agricultural activities. Approximately 17% of the total cultivated land in China is contaminated by Cd. Soil Cd pollution has become a serious global issue (Li et al., 2018).

Maize is an important agricultural crop worldwide. It is the predominant cereal in terms of planting area and global production. Maize makes a critical contribution to world food security and it is the main staple food in Africa and South America (Li et al., 2019). Cadmium in the soil can be easily absorbed by maize roots and then accumulate in plants and grains. Therefore, elucidating the molecular mechanism underlying Cd accumulation in maize has important implications for breeding maize varieties that accumulate low Cd levels, which will be useful for minimizing the Cd intake by humans.

Plants have evolved a series of regulatory mechanisms that protect them from Cd stress, including the following: (1) expelling or preventing the uptake of Cd by cells to minimize Cd accumulation (Cao et al., 2019; Yan et al., 2019; Zhang et al., 2019; Zhao et al., 2018a); (2) decreasing the Cd concentration in cells via chelation and complexation (Luo et al., 2018; Zhang et al., 2018c); and (3) increasing antioxidant enzyme activities to eliminate reactive oxygen species in cells (Gill and Tuteja, 2011; Rizwan et al., 2016). Among the identified Cd tolerance-related genes, those encoding transporter proteins have critical effects on the Cd accumulation in plants (Chen et al., 2019). The P1B-type heavy metal ATPases (HMAs), belonging to the P-type ATPase superfamily, are important for Cd transmembrane transport (Chen et al., 2019; Pedersen et al., 2012; Qiao et al., 2018; Takahashi et al., 2012a). Previous studies revealed that functionally diverse *HMA* genes encode proteins that vary regarding subcellular localization and tissue distribution. In *Arabidopsis thaliana* (Arabidopsis), AtHMA2 and AtHMA4 are localized in the plasma membrane and can promote Cd efflux from cells (Verret et al., 2004). Both *AtHMA2* and *AtHMA4* are primarily expressed in tissues surrounding the root vascular vessels and affect Cd root-to-shoot translocation (Verret et al., 2004). Additionally, AtHMA3 is localized in the vacuolar membrane, where it is responsible for detoxifying Cd through vacuolar sequestration (Morel et al., 2009). The *AtHMA3* gene is highly expressed in various tissues and cells, including the root apex, vascular tissues, guard cells, and hydathodes. The overexpression of *AtHMA3* can induce Cd accumulation in Arabidopsis shoots and roots (Morel et al., 2009). Similarly, OsHMA2 in the rice plasma membrane transports Cd out of cells (Takahashi et al., 2012b), whereas OsHMA3 functions in the vacuolar membrane to help sequester Cd in vacuoles (Miyadate et al., 2011; Ueno et al., 2010; Zhang et al., 2020). Although some genes regulating Cd accumulation have been functionally characterized in Arabidopsis and rice, the molecular mechanism regulating plant Cd tolerance remains unclear, and the key genes controlling Cd accumulation in maize have not been cloned.

The Cd accumulation level reportedly varies significantly among maize lines with differing genetic backgrounds (Zhao et al., 2018a). Applying molecular breeding technology to select and develop maize varieties that minimally accumulate Cd is an economically feasible and effective way to ensure the sustainable production of maize that is safe for human consumption (Jha and Bohra, 2016). In this study, we measured the maize grain Cd contents of an association panel comprising 513 lines. The results of a genome-wide association study (GWAS), bulked segregant RNA sequencing (BSR-seq), and fine mapping were combined to dissect the genetic loci controlling Cd accumulation in maize grains. By applying multiple approaches, including a gene functional annotation analysis, a sequence variation analysis, a transcript analysis, and an allelism test of the mutants, we revealed that a retrotransposon insertion in the *ZmCd1* gene affects the accumulation of Cd in maize grains. Phylogenetic and subcellular localization analyses indicated that *ZmCd1* might encode a P1B-type HMA transporter. On the basis of the sequence differences in the *ZmCd1* promoter, intron 1, intron 4, and exon 5 between maize lines that accumulate Cd at high and low levels, four PCR-based molecular markers were developed and five *ZmCd1* haplotypes were detected in maize lines. The predicted maize grain Cd accumulation in widely used inbred lines and hybrids in China based on molecular markers was consistent with the measured Cd contents. Therefore, *ZmCd1* is an important new target gene related to maize grain Cd accumulation. Moreover, allele-specific PCR markers may be applicable for breeding maize lines with low Cd levels.

## Results

### Differences in the Cd contents of 513 maize accessions

We measured the Cd contents of dry mature maize grains from 513 inbred lines of an association population harvested in two planting seasons (spring and autumn) in two fields [Zhuzhou (ZZ) and Ningxiang (NX)]. Examinations of the soil in the NX and ZZ fields (0–20 cm depth) revealed that the soil Cd content in ZZ (2.01–2.24 mg/kg) was higher than that in NX (1.69–1.85 mg/kg), and the ZZ field (pH 5.68–6.08) was more acidic than the NX field (pH 7.51–8.14) (**Figure 1a, b**). The maize grain Cd contents were higher in ZZ than in NX (**Figure 1c**), possibly because of the higher soil Cd content and lower soil pH in ZZ than in NX (Ali et al., 2019). The average maize grain Cd contents were 0.1955 and 0.2436 mg/kg in ZZ in the spring (ZZC) and autumn (ZZQ), respectively, whereas they were 0.0690 and 0.0457 mg/kg in NX in the spring (NXC) and autumn (NXQ), respectively. A combined analysis of the data for the four environments using the best linear unbiased prediction (BLUP) method determined the average maize grain Cd content was 0.1355 mg/kg (**Figure 1c; Table 1**). The data for both locations and the BLUP data indicated the maize grain Cd content varied substantially in the association population (**Table 1; Figure S1**). For example, the maize grain Cd content based on the combined analysis of all four environments ranged from 0.0358 to 0.7325 mg/kg (**Table 1**). The Pearson correlation coefficient reflected a moderately strong relationship among the maize grain Cd contents of the four environments (*r* = 0.54–0.68) (**Figure 1d**). An analysis of variance (ANOVA) revealed significant variations in the maize grain Cd contents based on the genotype, environment, and the genotype × environment interaction. The broad-sense heritability for the maize grain Cd content was as high as 71.11% (**Table 2**).

**Figure 1.**
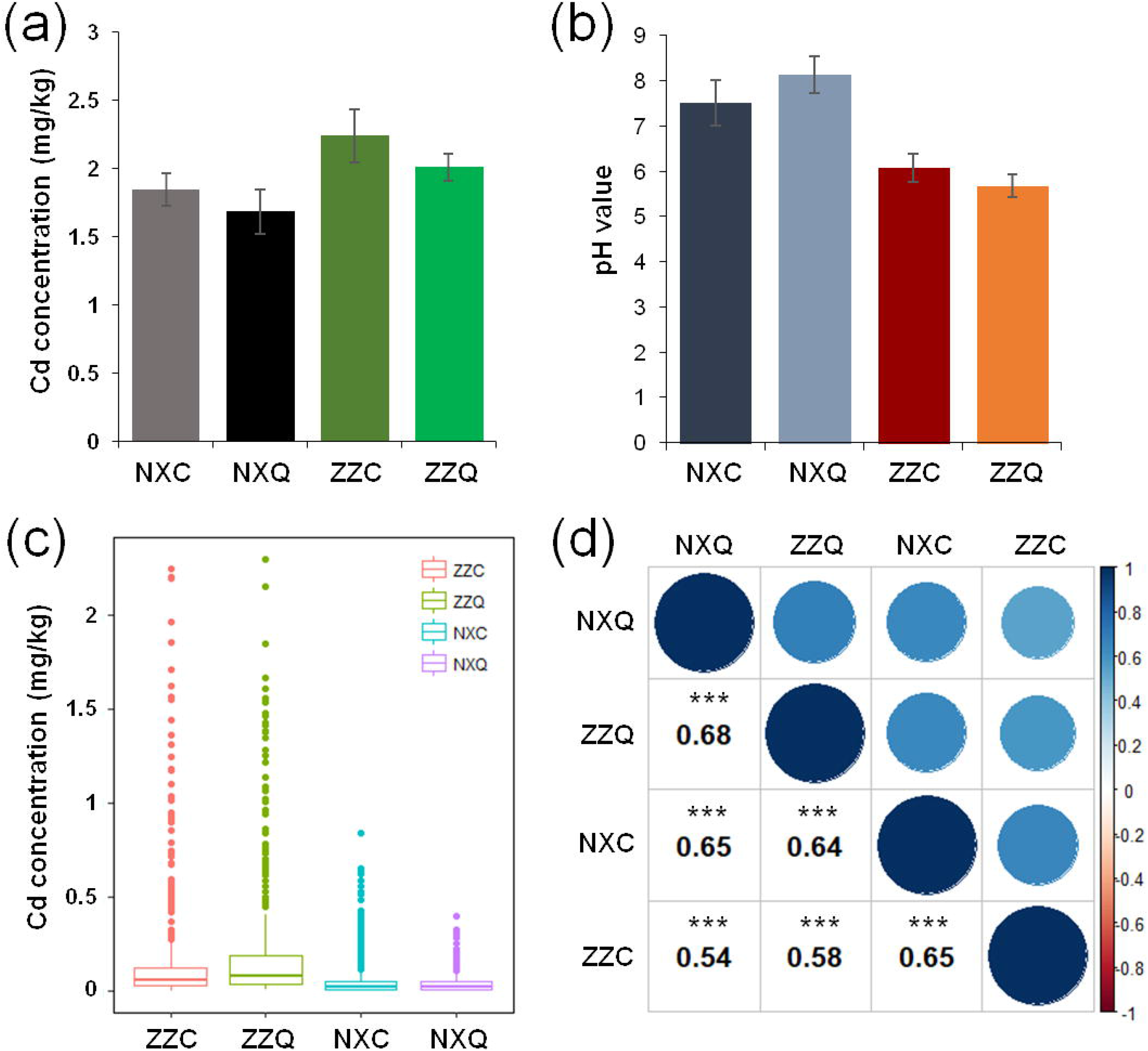
Maize grain Cd contents and correlations in the GWAS population in four environments. (a, b) Soil Cd content and pH in the spring and autumn in Zhuzhou (ZZ) and Ningxiang (NX) fields. Error bar = standard error, n = 5. (c, d) Maize grain Cd contents and correlations in the GWAS population in four environments. Correlation coefficients are provided in (d). Circle size represents the extent of the correlation. Blue and red indicate positive and negative correlations between traits, respectively. The color intensity is proportional to the extent of the correlation. ZZC, spring in ZZ. ZZQ, autumn in ZZ. NXC, spring in NX. NXQ, autumn in NX. \*\*\**P*< 0.001.

**Table 1.** Summary of the maize grain Cd contents of the GWAS population

| Location | Mean±SE (mg/kg) | Range (mg/kg) | Kurt | Skew |
| --- | --- | --- | --- | --- |
| NXC | 0.06904±0.005653 | 0.0002833-0.8412 | 11.68 | 3.152 |
| NXQ | 0.04571±0.002713 | 0.00078-0.4013 | 10.72 | 2.882 |
| ZZC | 0.1955±0.01859 | 0.00504-2.242 | 11.79 | 3.319 |
| ZZQ | 0.2436±0.02102 | 0.00754-2.298 | 6.816 | 2.603 |
| BLUPZZ | 0.2155±0.01056 | 0.06721-1.337 | 6.144 | 2.550 |
| BLUPNX | 0.05705±0.002473 | 0.01900-0.4168 | 11.21 | 3.078 |
| BLUP4EN | 0.1355±0.005328 | 0.03575-0.7325 | 6.341 | 2.532 |

**Table 2.** Analysis of variance for the Cd contents of the GWAS population in four environments

| Variance | MS | F | P value | H<sup>2</sup>(%) |
| --- | --- | --- | --- | --- |
| E | 6.8679 | 1083.4769 | 0.0000E+000 | 71.11 |
| G | 0.2409 | 38.0122 | 0.0000E+000 |  |
| G*E | 0.0953 | 15.0383 | 0.0000E+000 |  |

### Loci associated with maize grain Cd contents identified by a GWAS

A GWAS was performed using 1.25 M high-quality single nucleotide polymorphisms (SNPs) with a minor allele frequency greater than 0.05 and an association panel containing 513 maize inbred lines (Yang et al., 2014). A mixed linear model incorporating the population structure and relative kinship as well as the Cd concentration data for each environment were used. The BLUP data for the Cd concentration in two seasons at each location (BLUPZZ and BLUPNX) and the BLUP data for the Cd concentration in four environments (BLUP4EN) were also used for the GWAS (**Figure 2a, b; Figure S2**).

**Figure 2.**
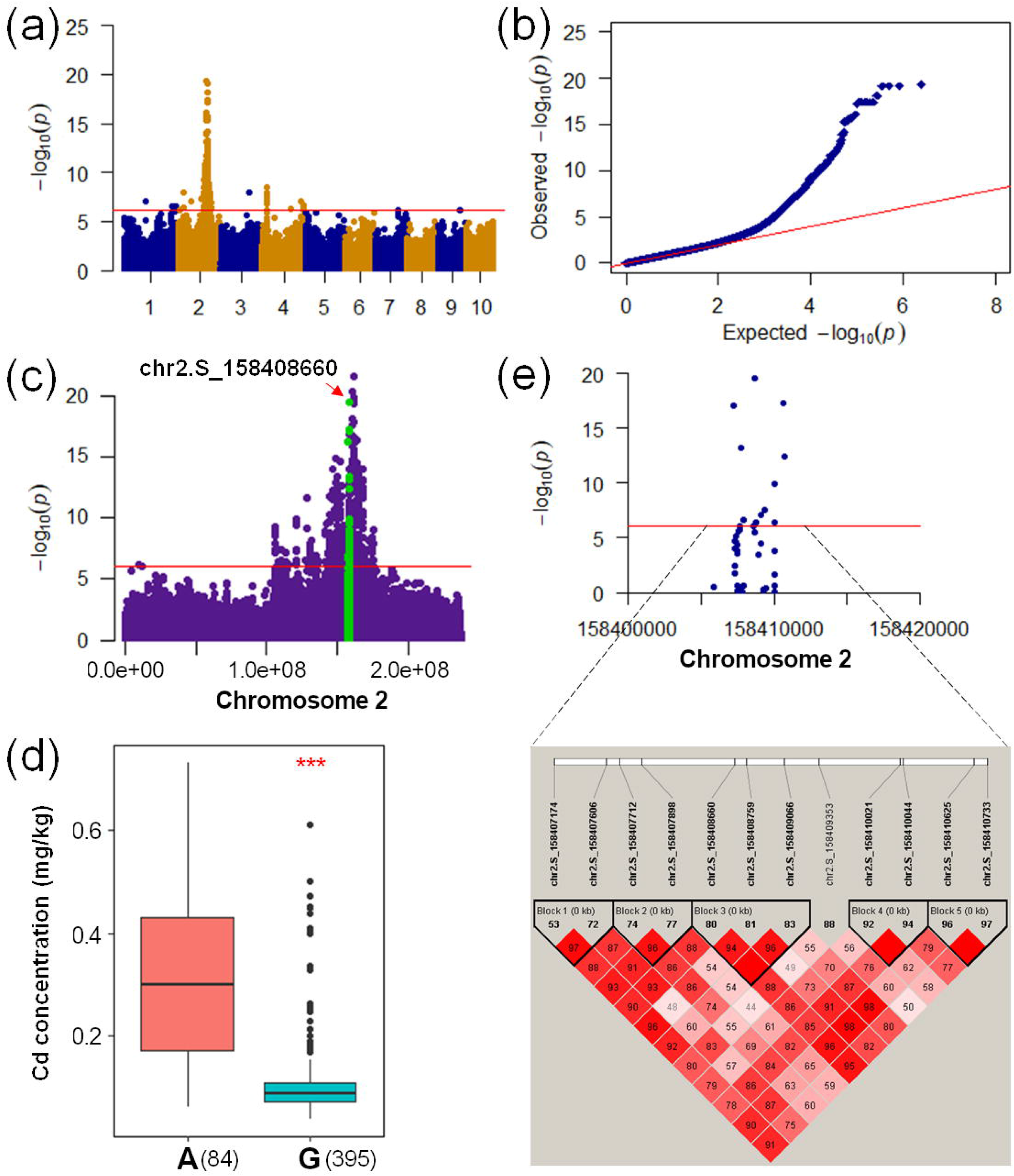
GWAS of the maize grain Cd accumulation. (a) Manhattan plots for the BLUP data of grain Cd accumulation across two seasons in NX. The red horizontal line represents the significance cutoff (*P* = 7.97e-7). NX, Ningxiang. (b) Quantile-Quantile plots for the GWAS MLM + Q + K model. (c) The GWAS signals for grain Cd accumulation are presented on chromosome 2. The peak SNP (i.e., SNP with the lowest *P*-value for the identified loci) for candidate gene *GRMZM2G175576* is indicated by a red arrow, whereas the SNPs within the identified QTL interval for grain Cd accumulation are highlighted in green. (d) Phenotypic differences resulting from the two alleles of the lead SNP (Chr2.S_158408660) of *GRMZM2G175576*. (e) Association between the *GRMZM2G175576* genetic variation and the grain Cd content as well as the linkage disequilibrium (LD) between the SNPs in *GRMZM2G175576*. The extent of the LD was determined based on the *D*′ statistic. Red squares without numbers represent complete LD (*D*′ = 1, *P* < 0.01). *D*′ values are presented in the squares. The red horizontal line represents the significance cutoff (*P* = 7.97e-7).

The linkage disequilibrium (LD) decay for the association population was 100 kb (r^2^ = 0.1) (Liu et al., 2017). Accordingly, the 100-kb region flanking both sides of each significant SNP was defined as a quantitative trait locus (QTL). A total of 55, 44, 34, and 71 QTL related to Cd accumulation were identified (*P* ≤ 7.97e-7) for ZZC, ZZQ, NXC, and NXQ, respectively (**Figure S2; Table S1**). Additionally, 68, 52, and 63 QTL were identified in the GWAS of BLUP4EN, BLUPNX, and BLUPZZ, respectively (**Figure S2; Table S1**). The phenotypic variance explained (PVE) for the QTL ranged from 5.24% to 23.78%. Moreover, 84.14% of the identified QTL were located on chromosome 2, with a PVE of 5.24%–23.78%. The remaining QTL were on eight other chromosomes, with a PVE of 5.43%–12.04% (**Table S1**). The QTL on chromosome 2 spanned a region from 5,468,732 bp to 189,546,923 bp (B73 RefGen_v2 reference genome). The PVE for these QTL gradually increased along the chromosome, but decreased after peaking. The QTL with the highest PVE were between 157,504,643 bp and 158,846,650 bp (PVE of 20.03%) and between 160,628,736 bp and 163,940,592 bp (PVE of 23.78%) on chromosome 2. They were detected simultaneously in all environments and in the BLUP data. Therefore, these two genomic regions harbor two major loci related to maize grain Cd accumulation, *qCd1* (peak SNP: chr2.S_158408660) and *qCd2* (peak SNP: chr2.S_161122814).

Three candidate genes (*GRMZM2G175576*, *GRMZM2G455491*, and *GRMZM2G018241*) in the two major QTL were proposed based on gene functional annotations (**Table 3**). Both *GRMZM2G175576* (*ZmHMA3*) and *GRMZM2G455491* (*ZmHMA4*) were detected in *qCd1*. These genes are homologous to Arabidopsis *HMA3* and rice *HMA3* genes, which are involved in sequestering Cd in root cell vacuoles. The other candidate gene, *GRMZM2G018241* (*ZmCESA9*), was detected in *qCd2* and encodes a cellulose synthase.

**Table 3.** Candidate genes for the maize grain Cd content revealed by a GWAS

| Traits | Peak SNP | Allele | Chr | QTL<br>interval <sup>a</sup> | *P*<br>value | *R*<sup>2</sup><br>(%) | Candidate<br>gene | Position | Annotation |
| --- | --- | --- | --- | --- | --- | --- | --- | --- | --- |
| BLUP4EN,<br>BLUPNX,<br>BLUPZZ,<br>NXC, NXQ,<br>ZZC, ZZQ | chr2.S_158408660 | G/A | 2 | 157504643-158846650 | 4.76E<br>-20 | 20.<br>03 | GRMZM2<br>G175576 | 158406991-158410957 | Heavy Metal-Transporting<br>P<sub>1B</sub>-Type ATPase |
| BLUP4EN,<br>BLUPNX,<br>BLUPZZ,<br>NXC, NXQ,<br>ZZC, ZZQ | chr2.S_158408660 | G/A | 2 | 157504643-158846650 | 4.76E<br>-20 | 20.<br>03 | GRMZM2<br>G455491 | 158386097-158389415 | Heavy Metal-Transporting P<sub>1B</sub>-Type<br>ATPase |
| BLUP4EN,<br>BLUPNX,<br>BLUPZZ,<br>NXC, NXQ,<br>ZZC, ZZQ | chr2.S_161122814 | G/A | 2 | 160628736-163940592 | 3.28E<br>-16 | 23.<br>78 | GRMZM2<br>G018241 | 161123950-161130108 | Cellulose synthase-9 |

<sup>a</sup>QTL intervals and candidate gene positions are based on the B73 AGPv2 genome sequence.
The QTL intervals comprise the 200-kb sequence ( ± 100 kb) flanking the peak SNP.

### A recessive gene was responsible for high maize grain Cd contents

Among the GWAS population inbred lines, Mo17 (0.4383 mg/kg) and Qi319 (0.4892 mg/kg) had high grain Cd levels, in contrast to the low grain Cd contents of B73 (0.0248 mg/kg) and Huangzaosi (0.0151 mg/kg) (**Figure 3a**). Maize inbred lines Jing724 (low Cd content; 0.0299 mg/kg) and D9H (high Cd content; 0.1283 mg/kg) were derived from maize hybrid x1132x, and they have been widely used for hybrid seed production in China. Grains from lines produced by crossing high-Cd maize lines (Mo17 and Qi319) (female parent) with low-Cd maize lines accumulated large amounts of Cd. Grains from lines generated by crossing low-Cd maize lines (B73, Huangzaosi, and Jing724) with high-Cd maize lines had low Cd levels (**Figure 3b**). These results suggested there was no pollen effect (xenia) on maize grain Cd accumulation.

**Figure 3.**
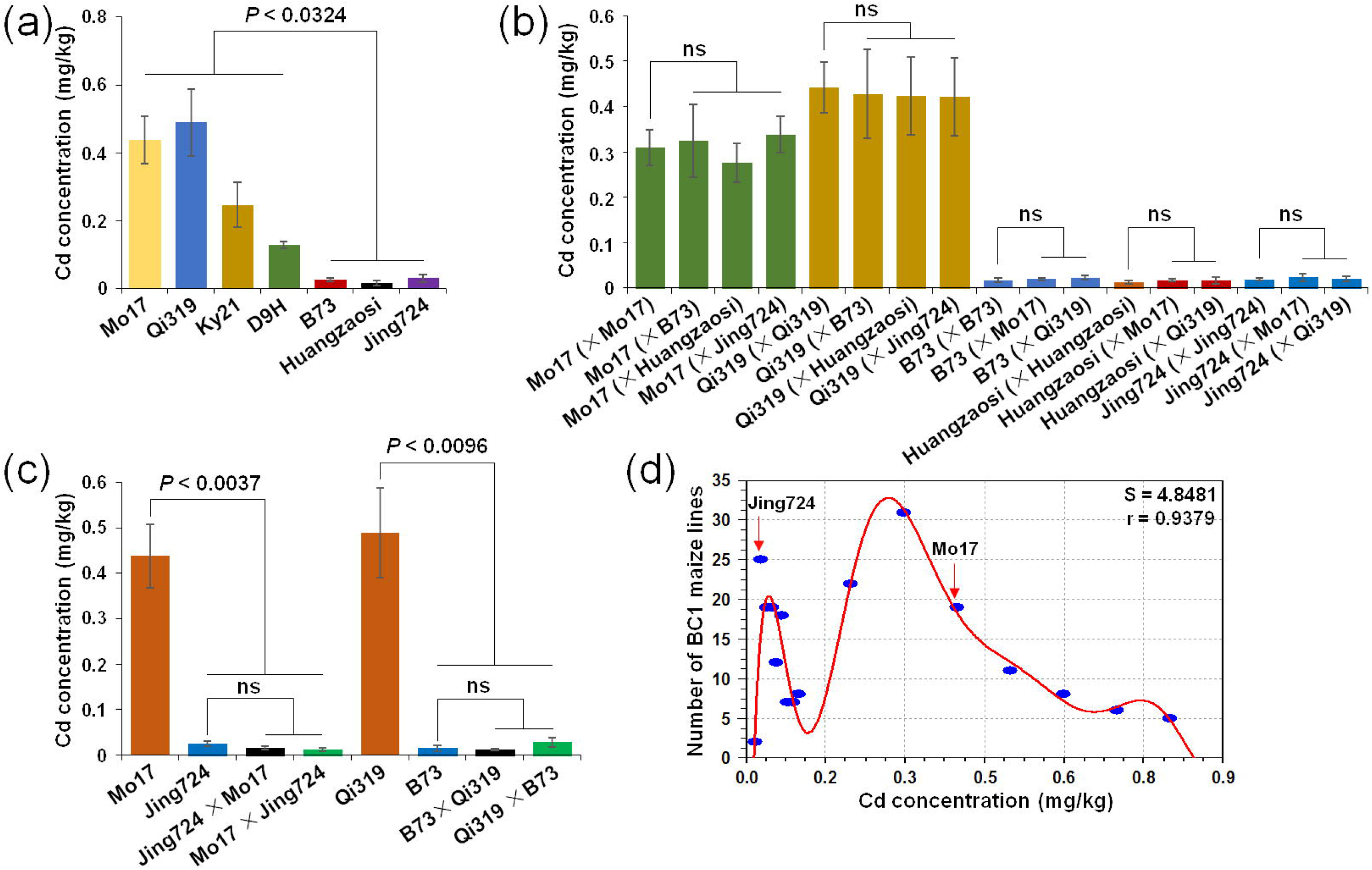
Grain Cd contents of maize inbred lines, hybrids, and the BC_1_ population of Jing724 and Mo17. (a) Grain Cd contents of seven maize inbred lines. (b) The Cd contents of heterozygous kernels were unaffected by foreign pollen grains. High-Cd lines (Mo17 and Qi319) were pollinated with the pollen grains of low-Cd lines (B73, Jing724, and Huangzaosi), resulting in heterozygous kernels with high Cd contents. The low-Cd lines were pollinated with the pollen grains from high-Cd lines, resulting in heterozygous kernels with low Cd contents. The inbred lines that provided the pollen grains are in parentheses. (c) Dominance of low grain Cd contents following the selfing of F1 hybrid plants. Hybrids were obtained by crossing the high-Cd inbred lines (Mo17 and Qi319) with the low-Cd inbred lines (Jing724 and B73). (d) Frequency distribution of the grain Cd contents of 218 plants in the BC_1_ population (Jing724 × Mo17) × Mo17. The curve generated by CurveExpert (version 1.4) (http://www.curveexpert.net/) revealed two apparent Cd content peaks using 0.1 mg/kg as the threshold: low Cd accumulation peak (n = 116) and high Cd accumulation peak (n = 102). Data were analyzed by Student’s *t*-test, n = 3 (a–c).

To analyze the effects of genes regulating Cd accumulation, Mo17 and Qi319 (high-Cd lines) were crossed with Jing724 and B73 (low-Cd lines). The grain Cd contents of the self-pollinated hybrids (Jing724 × Mo17, Mo17 × Jing724, B73 × Qi319, and Qi319 × B73) were low (0.0158, 0.0122, 0.0124, and 0.0285 mg/kg, respectively) (**Figure 3c**). Moreover, the Jing724 × Mo17 hybrid was backcrossed with Mo17 (high-Cd line) to generate a BC_1_ segregating population. The frequency distribution of the grain Cd content in the 218 BC_1_ plants revealed two apparent peaks, with a Cd content of 0.1 mg/kg set as the threshold (**Figure 4d**). Of the 218 BC_1_ plants, 102 had high grain Cd contents (0.1015–0.7678 mg/kg), whereas 116 had low grain Cd contents (0.0178–0.0938 mg/kg), which was consistent with the expected 1:1 ratio (χ^2^_1:3_ = 0.899, *P* = 0.3430) (**Figure 4d**). These results implied that the high maize grain Cd content was controlled by a single recessive major-effect gene.

**Figure 4.**
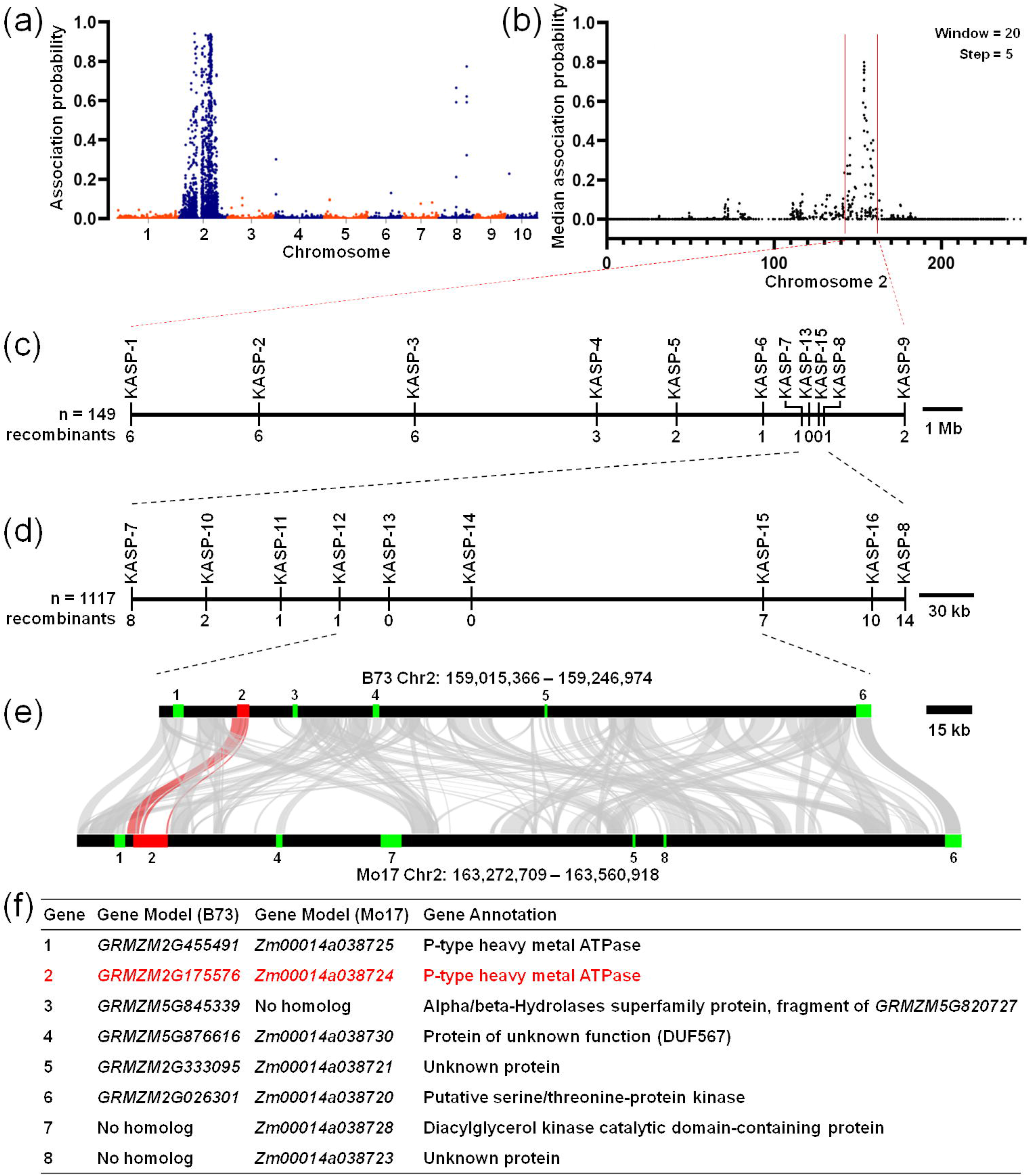
BSR-seq and fine mapping of the major QTL for the high grain Cd content in Mo17. (a) Probability of the association between 110,085 SNPs and the grain Cd accumulation. A BSR-seq analysis was performed using the RNA of two extreme Cd content pools: 30 plants with a high grain Cd content and 30 plants with a low grain Cd content from the F2 segregating population obtained by crossing Jing724 with Mo17 followed by selfing. The B73 RefGen_v3 genome was used as the reference sequence (https://www.maizegdb.org/) for the BSR-seq analysis and the subsequent fine mapping. (b) Significantly associated SNPs on chromosome 2. The median association probabilities for the SNPs on chromosome 2 were calculated using a sliding window method, with window and step sizes of 20 and 5 SNPs, respectively. (c, d) Fine mapping of *qCd1*. The *qCd1* QTL was mapped to the interval between KASP-7 and KASP-8 using 149 BC_1_ plants with a high grain Cd content (more than 0.2 mg/kg). New SNP-based KASP markers were developed to narrow the QTL interval to a 232-kb region between KASP-12 and KASP-15 using 1,117 high-Cd BC_1_ plants. (e) Synteny of the QTL intervals within the B73 and Mo17 genomes. Similar regions in the B73 and Mo17 sequences determined by a local BLAST analysis (e-value of 1e-100) are linked. Blue and red boxes represent the genes annotated in the corresponding regions of the B73 and Mo17 genomes. (f) Genes in the QTL intervals annotated in the B73 and Mo17 reference genomes. The candidate gene *GRMZM2G175576* is highlight in red (e, f).

### Mapping of the genes regulating Cd accumulation via BSR-seq

To produce extremely high and low Cd accumulation sample pools, an F2 segregating population was obtained by crossing Jing724 (low-Cd line) with Mo17 (high-Cd line). Total RNA samples extracted from the leaves of the high- and low-Cd plants in the F_2_ population were divided into two separate pools for an RNA sequencing (RNA-seq) analysis using the HiSeq X instrument (Illumina, USA). A total of 158 million reads were produced, with an average length of 2 × 150 bp/read. The raw reads were trimmed and then aligned to the B73 RefGen_v3 reference genome using the Genomic Short-read Nucleotide Alignment Program (GSNAP) (Wu et al., 2016). Reads with a single unique alignment were used for SNP calling.

To map the gene responsible for the Cd accumulation in grains, the high-Cd grains were used as the recessive pool, whereas the low-Cd grains were used as the dominant pool. A total of 110,085 SNPs were identified. The empirical Bayesian approach was employed to estimate the probability of a linkage between a high confidence SNP and the causal gene (Liu et al., 2012). The BSR-seq-based mapping revealed 946 associated SNPs clustered on chromosome 2 (**Figure 4a; Table S2**). To delimit the interval, a sliding window scan was performed for the associated SNPs (window, 20 SNPs; step, 5 SNPs), after which the significant SNPs were localized to the 142–161 Mb region on chromosome 2 (B73 RefGen_v3) (**Figure 4b**). The *qCd1* major QTL was located within this interval, suggesting it may be related to the grain Cd content variations between Jing724 and Mo17.

### Fine mapping of *qCd1* for maize grain Cd accumulation

To finely map *qCd1*, 11 SNP-based KASP (Kompetitive Allele-Specific PCR) markers (**Table S3**) were developed in the 142–161 Mb region on chromosome 2 for an analysis of 149 BC_1_ [(Jing724 × Mo17) × Mo17] individuals with high grain Cd contents (> 0.2 mg/kg). We mapped *qCd1* to a 423-kb interval between markers KASP-7 and KASP-8 (**Figure 4c**). Using another five SNP markers and an additional 1,117 BC_1_ individuals with high grain Cd contents (> 0.2 mg/kg), we further anchored *qCd1* to a 232-kb interval between markers KASP-12 and KASP-15 (**Figure 4d**). Six and seven genes were identified in this region in B73 and Mo17, respectively, based on the maize B73 RefGen_v3 reference genome (**Figure 4e, f**). Gene functional annotations indicated *GRMZM2G175576* (*ZmHMA3*) and *GRMZM2G455491* (*ZmHMA4*) encode P-type HMA transporters. Both genes were considered to be candidate genes in *qCd1*.

### *GRMZM2G175576* was identified as the candidate gene for maize grain Cd accumulation

To further characterize the candidate genes, the *GRMZM2G175576* and *GRMZM2G455491* genomic sequences and their expression were analyzed. The two gene sequences were identical in B73 and Jing724. Thus, B73 was used for further study because of the availability of the B73 reference genome. A comparison of the genomic sequences in Mo17 (high-Cd line) and B73 (low-Cd line) revealed 19 SNPs in the *GRMZM2G455491* coding sequence (CDS) that did not lead to translational termination. Regarding the comparison of the *GRMZM2G175576* sequences in Mo17 and B73, a 7,520-bp deletion and a 603-bp insertion were detected 300 bp upstream of the translation start codon as well as a 7,191-bp transposon insertion in intron 1 and a 628-bp insertion in intron 3 in Mo17 (**Figure 5a**). An analysis using the CENSOR software indicated the 603-bp insertion upstream of *GRMZM2G175576* was a gypsy-type LTR retrotransposon fragment, whereas the 7,191-bp insertion in exon 1 was a copia-type LTR retrotransposon and the 628-bp insertion in intron 3 was a LINE-1 retrotransposon.

**Figure 5.**
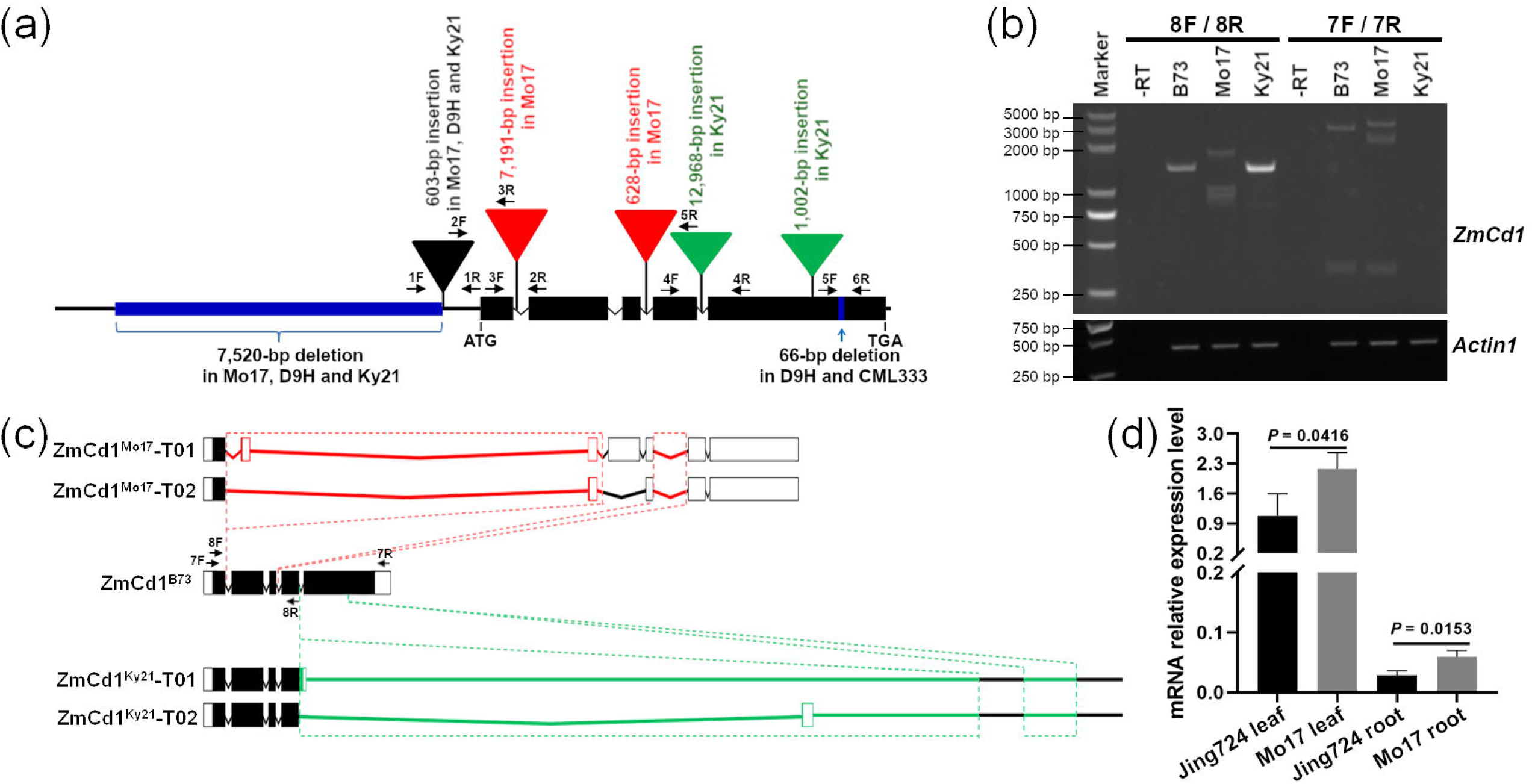
*ZmCd1* gene structural variations and expression. (a) *ZmCd1* genomic sequence variations in representative maize inbred lines. The backbone of gene structure was drawn based on the *ZmCd1^B73^* CDS and genomic sequences. Black boxes indicate exons, whereas black broken lines indicate introns. Insertions in the *ZmCd1* promoter, introns, and exons are labeled with triangles, whereas deletions in the promoter and exon 5 are marked in blue. (b) Results for the RT-PCR analysis of *ZmCd1* expression in B73, Mo17, and Ky21. The cDNA samples synthesized from the leaf RNA of B73, Mo17, and Ky21 seedlings were used for the PCR amplifications with primer pairs 7F/7R and 8F/8R. Primers 7F and 8F target exon 1, whereas primers 7R and 8R target exons 4 and 5, respectively, as shown in (c). Maize *Actin1* cDNA (*GRMZM2G126010*) was amplified as a control. The primers are listed in **Table S4**. The PCR products were analyzed by 2% agarose gel electrophoresis. M, DL5000 DNA Marker (Beijing TsingKe Biotech Co., Ltd, China). -RT, cDNA synthesis without a reverse transcriptase. (c) *ZmCd1* gene structures in B73, Mo17, and Ky21 determined by aligning DNA and transcript sequences. Boxes indicate exons, black boxes indicate coding regions, and black hat lines indicate introns. Only two representative *ZmCd1^Mo17^* transcripts are presented (**Figure S5**). The transposable element insertion regions within *ZmCd1^Mo17^* and *ZmCd1^Ky21^* genes are highlighted in red and green, respectively. (d) *ZmCd1* expression in Jing724 and Mo17 seedlings. Root and leaf total RNA samples for Jing724 and Mo17 at the V3 seedling stage were used for the qRT-PCR using primers targeting exon 1 (**Table S4**). Three biological replicates were used to calculate relative expression levels according to the 2^-ΔΔCt^ method. The *Actin1* gene served as the internal control. Data were analyzed by Student’s *t*-test. Error bar = standard error.

The data in the Maize eFP browser (http://bar.utoronto.ca/efp_maize/cgi-bin/efpWeb.cgi) suggested that *GRMZM2G175576* was highly expressed in the roots, leaves, internodes, and whole seeds, whereas *GRMZM2G455491* was highly expressed exclusively in the anthers (**Figure S3**). Reverse transcription polymerase chain reaction (RT-PCR) and quantitative real-time polymerase chain reaction (qRT-PCR) experiments revealed that *GRMZM2G175576* was expressed in maize seedling root and leaf tissues, in contrast to the lack of *GRMZM2G455491* expression in Jing724, B73 and Mo17 **(Figure S4)**. These results implied *GRMZM2G175576* was the more likely target gene.

The GWAS analysis indicated *GRMZM2G175576* harbors 12 significant SNPs, with the lead SNP (chr2.S_158408660, *P* = 4.76e-20) explaining 20.03% of the phenotypic variation (**Figure 2c**). The phenotypic differences associated with the two alleles of chr2.S_158408660 were significant (**Figure 2d**). The 12 significant SNPs in *GRMZM2G175576* were clustered (*D*′ > 0.94), reflecting the strong LD of these SNPs (**Figure 2e**). This result further confirmed that *GRMZM2G175576* was the candidate gene. Hereafter, we refer to *GRMZM2G175576* as *ZmCd1*.

### *ZmCd1* transcriptional variation between Mo17 and B73

A qRT-PCR analysis proved that *ZmCd1* was expressed in the B73 and Mo17 maize seedling leaf and root tissues. The *ZmCd1* expression level was significantly higher in Mo17 than in Jing724 (**Figure 5d**), suggesting that the 7,520-bp deletion and transposon insertion in the *ZmCd1* promoter resulted in increased expression in Mo17. To map the *ZmCd1* transcription start site (TSS) in B73 and Mo17, a 5′-RACE was conducted using gene-specific primers targeting exon 1 (**Table S4**). The *ZmCd1^B73^* TSS was 191 or 89 bp upstream of the start codon, whereas the *ZmCd1^Mo17^* TSS was 274 bp upstream of the start codon. Thus, differences in the *ZmCd1* promoter did not influence translation initiation in Mo17.

Using 7F/7R primers to amplify *ZmCd1* from exon 1 to exon 5, we determined that the amplified Mo17 *ZmCd1* sequences were nonspecific and diffuse, in contrast to the distinct 2,737-bp amplified B73 *ZmCd1* sequence (**Figure 5b**). The amplified *ZmCd1^Mo17^* fragments were cloned and sequenced, resulting in the detection of 19 different transcripts in 109 clones, of which 72 clones had transcripts shorter than 2,737 bp and the other 37 clones had transcripts longer than 2,737 bp (**Figure 5c; Figure S5**). The transcript analysis revealed that the retrotransposon inserted in intron 1 resulted in transcript and amino acid coding abnormalities in *ZmCd1^Mo17^* (Dataset S1). Therefore, we proposed that the retrotransposon in intron 1 abolished the *ZmCd1* function in Mo17, leading to increased grain Cd accumulation.

### Functional verification of *ZmCd1* by an allelism test of the mutants

Two B73 EMS mutants, *zmcd1-1* (Mut_Sample: EMS4-038f45) and *zmcd1-2* (Mut_Sample: EMS4-038f39), were obtained from the maize EMS mutant library (http://www.elabcaas.cn/memd/) for an allelism test. Both *zmcd1-1* and *zmcd1-2* have a termination mutation in the *ZmCd1* coding region (**Figure 6a, b**). The Cd contents in the *zmcd1-1* and *zmcd1-2* maize grains harvested from Cd-contaminated farmland were as high as 0.3507 and 0.3093 mg/kg, respectively (**Figure 6c**). Both maize mutants were crossed with the high-Cd maize line Mo17. The Cd contents of the grains from the self-pollinated hybrid maize plants grown on Cd-contaminated farmland were determined. Grains from Mo17 × *zmcd1-1* and *zmcd1-2* × Mo17 self-pollinated lines had high Cd levels of 0.4008 and 0.3421 mg/kg, respectively (**Figure 6c**). The allelism test results verified that *ZmCd1* was associated with the high maize grain Cd content.

**Figure 6.**
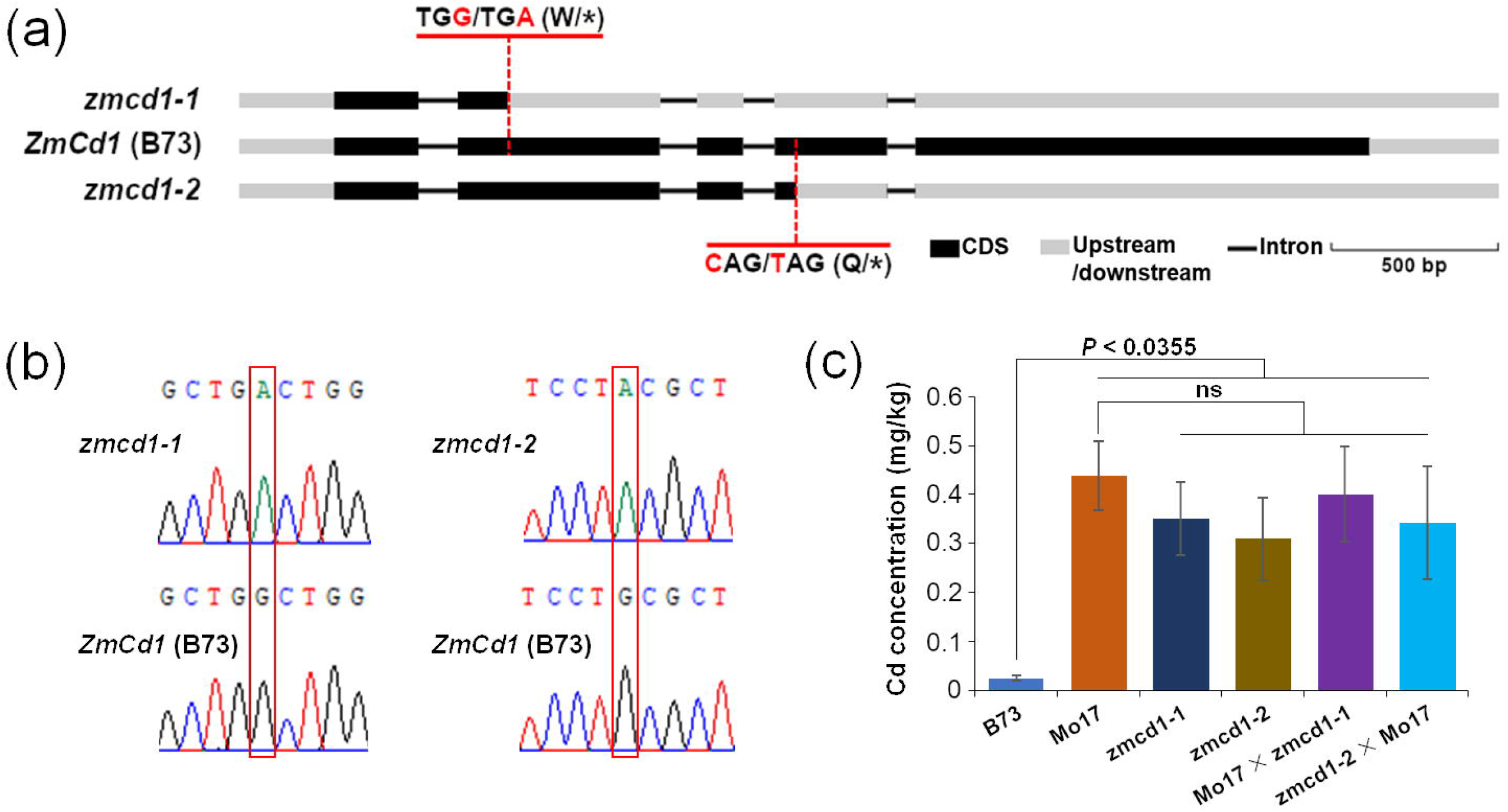
High grain Cd contents of *ZmCd1* EMS mutants and results of the allelism test. (a) *ZmCd1* gene mutation sites in the *zmcd1-1* (EMS4-038f45) and *zmcd1-2* (EMS4-038f39) EMS mutants. (b) Validation of mutation sites by DNA sequencing. Altered nucleotides in the EMS mutants are framed by red dashed lines. (c) Allelism test results for the *ZmCd1* mutants and Mo17. Maize plants grown in a Cd-contaminated field in Zhuzhou in the autumn of 2018 were harvested for grain Cd content measurements. The grain Cd contents were significantly higher for the *zmcd1-1* and *zmcd1-2* mutants than for B73 (wild-type). The kernels produced by the F1 hybrids, Mo17 × *zmcd1-1* and *zmcd1-2* × Mo17, had high Cd contents. Data were analyzed by Student’s *t*-test (n = 4). Error bar = standard error.

### Evolutionary analysis and subcellular localization of ZmCd1

A phylogenetic analysis divided 43 HMA-encoding genes from Arabidopsis, rice, sorghum, maize, *Brassica rapa*, barley, and durum wheat into four clades (**Figure 7a; Table S5**). Specifically, *ZmCd1/ZmHMA3* in the Zn/Co/Cd/Pb clade formed a subclade with *OsHMA3*, *SbHMA3*, *TdHMA3-B1a*, and *HvHMA3-B* (**Figure 7a**), indicating that *ZmCd1* is probably functionally similar to *OsHMA3* and *AtHMA3*. A protein sequence alignment revealed a 66.5% sequence match between *ZmCd1* and *OsHMA3* and a 51.9% sequence match between *ZmCd1* and *AtHMA3* (**Figure S6**).

**Figure 7.**
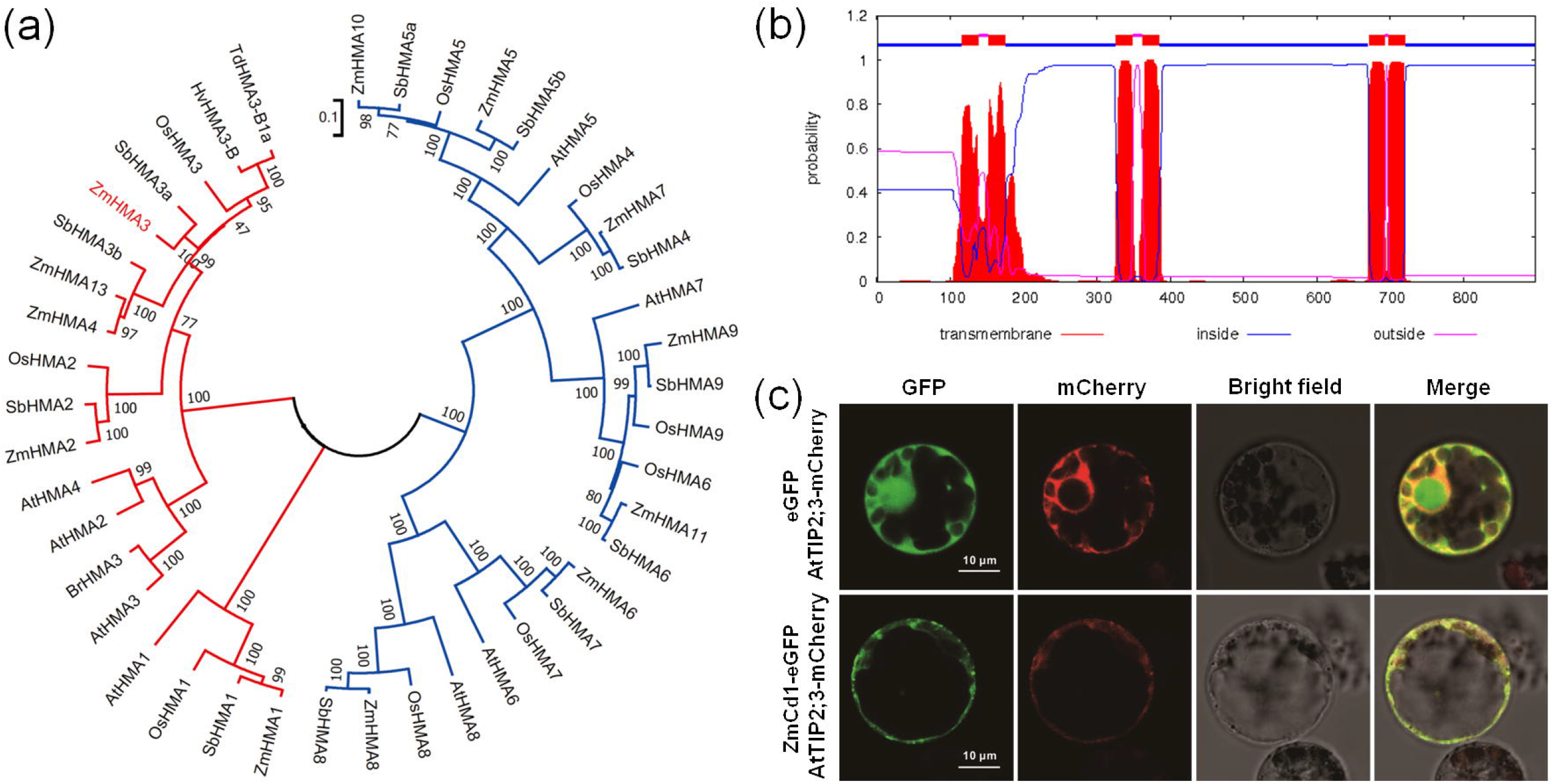
Phylogenetic relationships among plant HMAs and the subcellular localization of ZmCd1. (a) Phylogenetic tree of plant HMAs. The neighbor-joining tree was constructed using MEGA 7 based on the alignment of 43 plant HMA proteins, including 8 AtHMAs in Arabidopsis (Baxter et al., 2003), 9 OsHMAs in rice (Baxter et al., 2003; Zhiguo et al., 2018), 9 SbHMAs in sorghum (Zhiguo et al., 2018), 12 ZmHMAs in maize (Cao et al., 2019; Zhiguo et al., 2018), BrHMA3 in *Brassica rapa* (Zhang et al., 2019), HvHMA3-B in barley (Lei et al., 2020), and TdHMA3-B1a in durum wheat (Maccaferri et al., 2019). The Poisson amino acid substitution model and 1,000 bootstrap replicates were used. The *ZmCd1* (*ZmHMA3*) gene is highlighted in red. (b) ZmCd1 transmembrane domains predicted by the TMHMM server (version 2.0) (http://www.cbs.dtu.dk/services/TMHMM/). (c) Transient expression of the ZmCd1-eGFP fusion protein in maize protoplasts. The pM999-ZmCd1-eGFP and pM999-AtTIP2;3-mCherry recombinant plasmids were used for the co-transformation of maize protoplasts, which were then examined for fluorescence. The Arabidopsis TIP2;3 protein, which served as a tonoplast-localized control, was anchored to the vacuolar membrane (Zhang et al., 2020). Maize protoplasts co-transformed with pM999-eGFP (blank) and pM999-AtTIP2;3-mCherry were used as the negative control. Scale bar = 10 μm.

Because *ZmCd1* was predicted to encode six putative transmembrane domains (**Figure 7b**), we performed a maize protoplast transient expression assay using the ZmCd1-eGFP fusion protein and the protoplast-localized fusion protein comprising Arabidopsis TIP2;3 and mCherry. The ZmCd1-eGFP fusion protein was co-localized with AtTIP2;3-mCherry in the maize protoplast vacuolar membrane (**Figure 7c**). We speculated that ZmCd1 helps sequester Cd in vacuoles, similar to AtHMA3 and OsHMA3.

### Natural variations in *ZmCd1* in diverse maize lines

Using the structural variations of the *ZmCd1* gene in 38 maize inbred lines with sequenced genomes, six simple natural *ZmCd1* variation types (haplotypes) were identified based on the insertions/deletions (InDels) in the *ZmCd1* promoter, intron 1, intron 4, and exon 5 (compared with the B73 sequence) (**Table S6; Figure 5a**). The first variation type (i.e., InDels in the promoter) was detected only in the European EP1 and DK105 maize lines. The five other types of *ZmCd1* variations were detected in the maize lines examined in this study, and the representative maize lines were B73, Mo17, Ky21, D9H, and CML333, respectively (**Figure 5a; Figure 8**). Notably, D9H was used to represent SK, CML228, and F7 because the InDel variation in the *ZmCd1^D9H^* promoter was similar to that in the Mo17 *ZmCd1* promoter.

**Figure 8.**
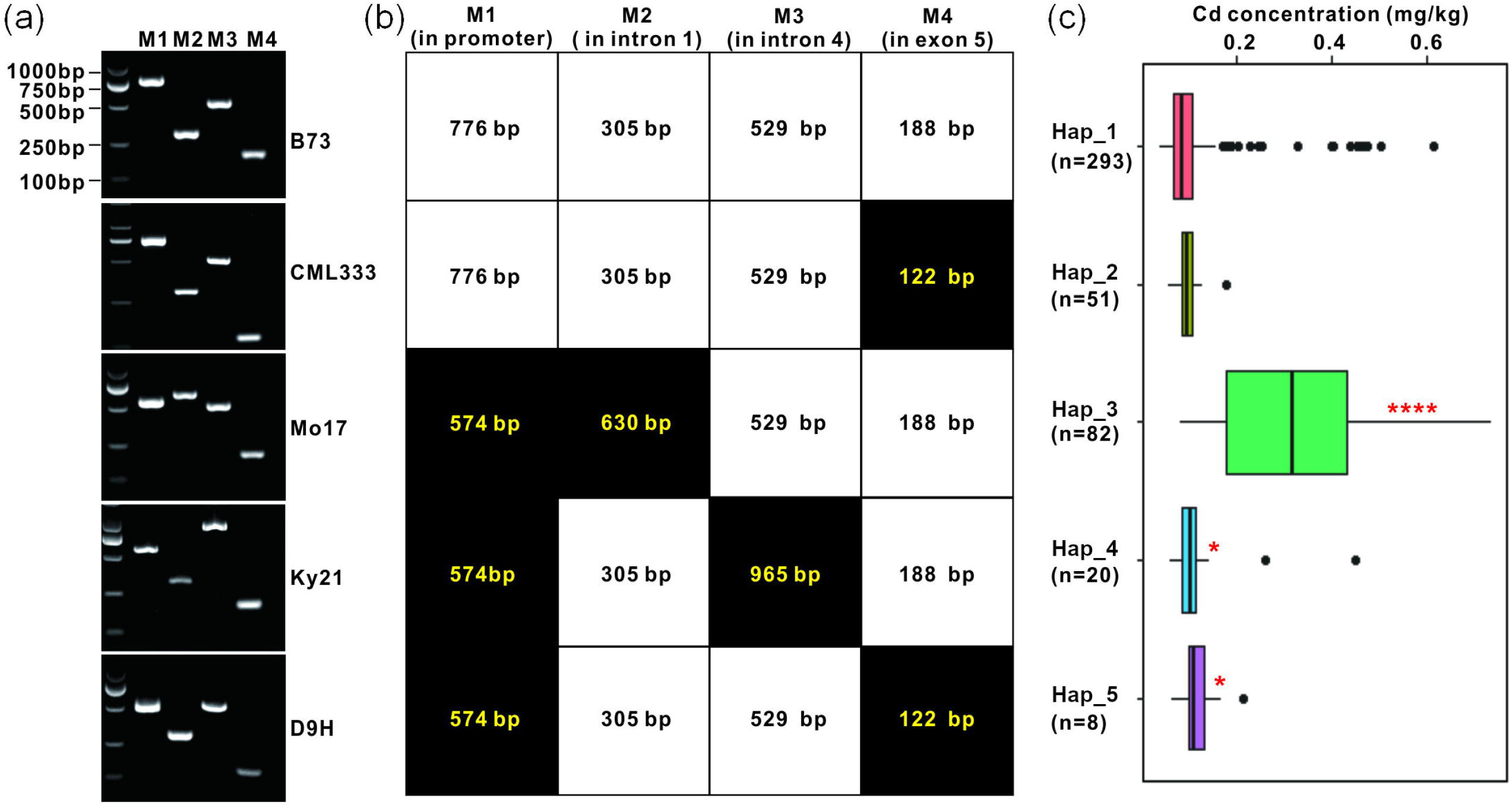
*ZmCd1* gene haplotypes in the GWAS panel. (a) Analysis of the PCR-based molecular markers by 2% agarose gel electrophoresis. The following primers for the M1–M4 molecular markers were designed based on the gene structural variations (**Figure 5a** and **Table S6**): 1F/2F//1R for M1, 3F//2R/3R for M2, 4F//4R/5R for M3, and 5F/6R for M4. Regarding M1–M3, three primers were used in each PCR amplification and the PCR extension time was less than 1 min. The left lane presents the DL2000 DNA Marker (Beijing TsingKe Biotech Co., Ltd, China). The genomic DNA from five representative maize inbred lines (B73, CML333, Mo17, Ky21, and D9H) harboring five different haplotypes of the M1–M4 molecular marker combination was used for the PCR amplification. (b) The M1–M4 molecular marker lengths in B73, CML333, Mo17, Ky21, and D9H. Black boxes indicate the molecular markers that differed in B73. (c) Grain Cd contents for five haplotypes in the GWAS panel. Student’s *t*-test was used to compare haplotypes with hap_1. \**P*< 0.05, \*\*\**P*< 0.001.

Compared with the corresponding sequence in B73 (low-Cd line, 0.0248 mg/kg), *ZmCd1* in Ky21 (high-Cd line, 0.25 mg/kg) had a 7,520-bp deletion and a 603-bp insertion 300 bp upstream of the translation start codon as well as a 12,968-bp gypsy-type LTR retrotransposon insertion in exon 4 and a 1,002-bp MuDR DNA transposon insertion in exon 5 (**Figure 5a**). The full-length *ZmCd1* cDNA sequence was amplified by PCR using the 7F/7R primers, but no amplified fragment was detected for Ky21. However, a 1,355-bp fragment was detected when the 8F/8R primers were used to amplify the sequence between *ZmCd1* exons 1 and 4 (**Figure 5b**). These results indicated that the insertion mutation in intron 4 resulted in a premature transcriptional termination in Ky21. The 3′ ends of *ZmCd1^Ky21^* transcripts were obtained by 3′-RACE using gene-specific primers targeting exon 4. A sequence analysis revealed two transcripts that were shorter in Ky21 than in B73, with no exon 5. One transcript retained the 1–139 bp sequence of intron 4, whereas the other transcript retained the 9,601–9,816 bp sequence of intron 4 (**Figure 5c**).

Compared with the B73 *ZmCd1* sequence, the *ZmCd1* gene in maize inbred line D9H (high-Cd line, 0.13 mg/kg) had a 7,520-bp deletion and a 603-bp insertion 300 bp upstream of the translation initiation codon as well as a 66-bp deletion in exon 5. In maize inbred line CML333 (low-Cd line, 0.06 mg/kg), the *ZmCd1* sequence only had a 66-bp deletion in exon 5 (**Figure 5a**).

### *ZmCd1* molecular marker development

Four PCR-based molecular markers were developed based on the *ZmCd1* differences in the upstream (M1) sequence and in intron 1 (M2), intron 4 (M3), and exon 5 (M4) among B73, CML333, Mo17, Ky21, and D9H (**Figure 5a; Figure 8a**). Using these markers, five haplotypes were detected in the maize lines included in this study. In Hap_1, the PCR amplified *ZmCd1* products were 776, 305, 529, and 188 bp in B73 (low-Cd line). In Hap_2, the PCR amplified products were 776, 305, 529, and 122 bp in CML333 (low-Cd line). In Hap_3, the PCR amplified products were 574, 630, 529, and 188 bp in Mo17 (high-Cd line). In Hap_4, the PCR amplified products were 574, 305, 965, and 188 bp in Ky21 (high-Cd line). In Hap_5, the PCR amplified products were 574, 305, 529, and 122 bp in D9H (high-Cd line) (**Figure 8a, b**).

The genotyping of 454 maize lines from the GWAS population using the four molecular markers developed for *ZmCd1* indicated that 293 lines carried Hap_1, 51 lines carried Hap_2, 82 lines carried Hap_3, 20 lines carried Hap_4, and 8 lines carried Hap_5. An analysis of the BLUP data for the maize lines with haplotypes 1–5 determined that the average Cd contents were 0.0993, 0.0939, 0.3163, 0.1214 and 0.1199 mg/kg, respectively. Moreover, there was a significant difference in the Cd content between Hap_1 and Hap_3 (*P* < 0.0001), Hap_1 and Hap_4 (*P* < 0.05), and Hap_1 and Hap_5 (*P* < 0.05) (**Figure 8c**).

### Prediction of maize grain Cd accumulation using the molecular markers developed in this study

The genotypes of 36 elite inbred lines and 13 elite hybrids widely used for maize production were analyzed using the four developed markers. The 13 hybrids included the top three varieties in China (Xianyu335, Zhengdan958, and Jingke968). Four *ZmCd1* haplotypes were detected in the 36 inbred lines, including 20 lines carrying Hap_1, 7 lines carrying Hap_3, 3 lines carrying Hap_4, and 6 lines carrying Hap_5 (**Figure 9a**). We detected three haplotypes in 13 hybrids, with Zhengdan958 carrying Hap_1/Hap_3, Xianyu335, MC812, and Jingnongke736 carrying Hap_1/Hap_5, and Yuqingzhu23 and Kangnong999 carrying Hap_3 (**Figure 9b**). These maize inbred lines and hybrids were grown in Cd-contaminated soil, after which their grain Cd contents were measured. The maize grain Cd contents of these materials were consistent with the results predicted using the molecular markers (**Figure 9**). The grain Cd contents of 20 inbred lines (e.g., Jing2416, Jing724, MC01, and Jing92) and 7 hybrids (e.g., Jingnongke728 and Jingke968) carrying Hap_1 were low (0.0096–0.0577 mg/kg), in contrast to the high grain Cd contents (0.1094–1.6760 mg/kg) of 16 inbred lines carrying Hap_3, Hap_4, or Hap_5 and two hybrids carrying Hap_3. Hybrids Zhengdan958, MC812, Xianyu335, and Jingnongke736 carried Hap_1/Hap_3 or Hap_1/Hap_5. Because Hap_1 is dominant over Hap_3 and Hap_5, these four hybrids had low grain Cd contents (**Table S7**). These results proved that the four molecular markers can be used to predict maize grain Cd accumulation characteristics, making them useful for screening low-Cd maize germplasms and for improving high-Cd maize lines by molecular marker-assisted selection.

**Figure 9.**
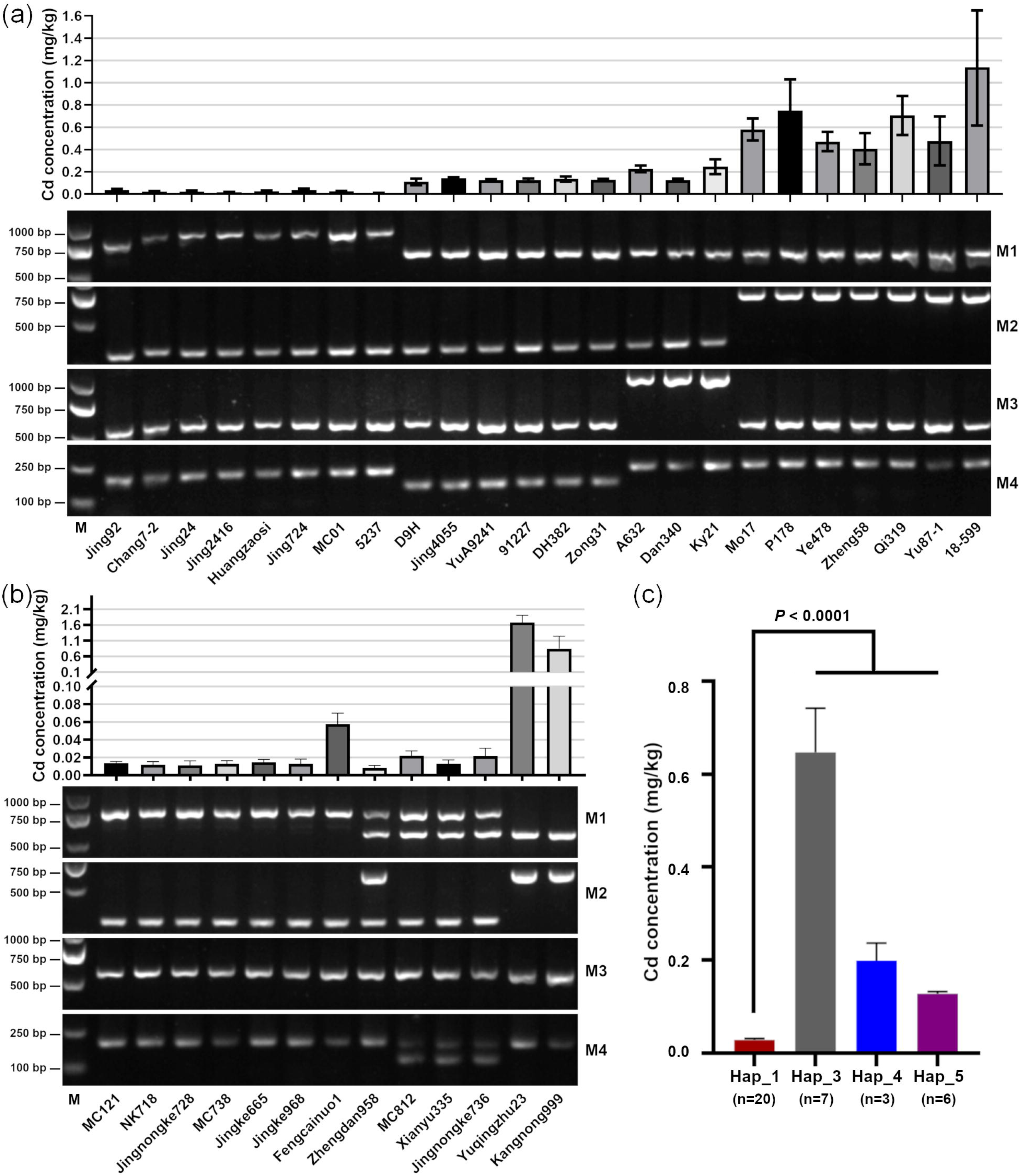
Application of PCR-based markers for evaluating the grain Cd contents of maize inbred lines and hybrid varieties. (a, b) Grain Cd contents (upper panel) and detection of the M1–M4 molecular markers (lower panel) for 24 maize inbred lines (a) and 13 maize hybrid varieties (b). Maize plants were grown in a Cd-contaminated field in Zhuzhou in the autumn of 2018 for an analysis of grain Cd contents. Details regarding the maize inbred lines and hybrid varieties are listed in **Table S7**. M, DL2000 DNA Marker (Beijing TsingKe Biotech Co., Ltd, China). (c) Grain Cd contents in inbred lines with differing haplotypes. The grain Cd contents are listed in **Table S7**. Data were analyzed by Student’s *t*-test. Error bar = standard error.

## Discussion

Minimizing maize grain Cd accumulation is critical for decreasing the risks of Cd toxicity in humans. Dissecting the genetic components underlying Cd accumulation may enable researchers and breeders to decrease maize grain Cd contents (Jha and Bohra, 2016; Liu et al., 2019). In this study, combining multiple genetic mapping techniques led to the identification of a candidate gene for maize grain Cd accumulation on chromosome 2. The allelism test of the mutants confirmed *ZmCd1* contributes to grain Cd accumulation. A DNA sequence analysis detected natural variations in the *ZmCd1* promoter, intron 1, intron 4, and exon 5. A transcript analysis revealed mutations in transcripts and the amino acid-encoding sequence of *ZmCd1* resulting from a retrotransposon insertion. Subsequent phylogenetic and subcellular localization analyses indicated that *ZmCd1* encodes a vacuolar membrane-localized P_1B_-type HMA transporter. Moreover, four molecular markers for *ZmCd1* were developed and five *ZmCd1* haplotypes were detected. The utility of the molecular markers for predicting the maize grain Cd accumulation phenotype confirmed that these molecular markers are useful for maize breeding.

Cadmium accumulation is related to environmental factors (Gill and Tuteja, 2011; Zhang et al., 2018a). Several studies have demonstrated that plant Cd contents are influenced by soil Cd concentrations, pH, and other factors (Ali et al., 2019; Florijn and Van Beusichem; Gill and Tuteja, 2011). In rice, Cd accumulation increases as the pH of the culture medium increases from 3.5 to 6.0, but it decreases as the pH increases to 7.0 and 8.0 (Ali et al., 2019). Similar to these findings, we determined that the average maize grain Cd content in two seasons in ZZ was higher than that in NX, which is consistent with the observation that the soil Cd concentration in ZZ was higher than that in NX. Additionally, the soil pH in ZZ was close to 6.0, whereas it was higher than 7.5 in NX (**Figure 1b**).

In this study, a large genotypic variation in the maize grain Cd content was detected for the GWAS population regardless of the season and location, suggesting that maize grain Cd accumulation is genetically controlled. Previous studies identified several factors regulating Cd tolerance, among which ATP-binding cassettes (Patrizia et al., 2015; Zhang et al., 2018b) and HMAs (Jha and Bohra, 2016; Maccaferri et al., 2019; Sasaki et al., 2014; Yan et al., 2016) have been widely investigated in Arabidopsis, rice, and wheat. However, the key gene regulating Cd accumulation in maize remained uncharacterized. In a recent study, a major QTL (153.75–167.58 Mb on chromosome 2) for maize leaf Cd accumulation, *qLCd2*, was identified based on a GWAS and a DH population (Zhao et al., 2018a). The genomic position of *qLCd2* was comparable to that of the major QTL identified by BSR-seq (142–161 Mb) and GWAS mapping in this study, suggesting that this QTL controls Cd accumulation in maize leaves and grains. The fine mapping results anchored this major QTL to the 159,015,366–159,246,974 bp interval on chromosome 2. Furthermore, gene functional annotations and sequence analyses resulted in the identification of *GRMZM2G175576* (*ZmCd1*) as a candidate gene. The *ZmCd1* function was further verified by an allelism test of the maize *zmcd1* mutants.

The *ZmCd1* sequence of the high-Cd maize line Mo17 included long DNA deletion and retrotransposon insertion variations in the promoter and introns. A previous investigation proved that the differences in *OsHMA3* expression and Cd accumulation between *japonica* and *indica* rice were due to 11 nucleotide differences in the promoter (Liu et al., 2020). A Sukkula-like transposable element insertion upstream of *HvHMA3* reportedly promotes gene expression (Lei et al., 2020). Similarly, we observed that *ZmCd1* expression was significantly higher in Mo17 than in B73 (**Figure 5d**), possibly because of the insertion of a 603-bp retrotransposon in the promoter. Although the *ZmCd1* expression level was higher in Mo17, a 7,191-bp retrotransposon insertion in intron 1 resulted in mutated transcripts (**Figure 5a–c**) with abnormal amino acid-encoding sequences. The resulting loss of functional ZmCd1 increased the grain Cd content.

A phylogenetic analysis of maize, barley, durum wheat, sorghum, *B. rapa*, Arabidopsis, and rice revealed that *ZmCd1* belongs to the Zn/Co/CD/Pb group and forms a subclade with *OsHMA3*, *SbHMA3*, *TdHMA3-B1a*, and *HvHMA3-B*. The OsHMA3, HvHMA3-B, and TdHMA3-B1a proteins are localized to the vacuolar membrane (Lei et al., 2020; Maccaferri et al., 2019; Miyadate et al., 2011), where they help sequester Cd in vacuoles and influence root-to-shoot Cd translocation (Lei et al., 2020; Maccaferri et al., 2019; Miyadate et al., 2011; Ueno et al., 2010). These findings suggest the HMA3 transporters mediate a conserved Cd tolerance mechanism in diverse species. In this study, a subcellular localization analysis confirmed that ZmCd1 is a vacuolar membrane protein (**Figure 7c**). The hybrid grain Cd contents were correlated with the corresponding Cd content of the female parent (**Figure 3b**), indicative of a lack of xenia effect on the maize grain Cd content. Consistent with the distribution of Cd in various rice tissues (Rizwan et al., 2016), the rank order of the Cd contents in maize tissues was as follows: leaf > stem > grain (**Figure S7**). These results indicate that ZmCd1 mediates the sequestration of Cd in vacuoles and restricts the Cd accumulation in maize grains by modulating Cd root-to-shoot translocation.

We developed four molecular markers based on the natural variations in *ZmCd1* among 38 maize inbred lines with sequenced genomes. Five variation types (Hap_1–5) were identified in the study materials, and three (Hap_3–5) were associated with high maize grain Cd contents. The *ZmCd1* genes of the Hap_3 and Hap_4 lines had a transposon inserted in introns, which adversely affected transcription. The Hap_5 lines had a 66-bp deletion in *ZmCd1* exon 5. These three haplotypes had abnormal amino acid-encoding *ZmCd1* sequences, resulting in nonfunctional ZmCd1 (**Dataset S1**). However, the Hap_2 and Hap_5 both having the 66-bp deletion in exon 5 showed different grain Cd contents probably due to the presence of other loci in the GWAS panel controlling Cd accumulation in maize grains, which would be figured out in the further study. The analysis of the GWAS population indicated most of the maize inbred lines had low Cd levels. Additionally, Hap_3 was the main haplotype among the high-Cd maize lines. The molecular markers were used to investigate the commonly cultivated maize inbred lines and hybrids in China. Hybrids Jingke968, Zhengdan958, and Xianyu335 are the predominant maize varieties grown in China. Jingnongke728 (MC01 × Jing2416) is suitable for mechanical harvesting. The Cd accumulation of 13 hybrids (including the above-mentioned hybrids) and 36 inbred lines was consistent with that predicted by the molecular markers (**Figure 9; Table S7**), implying the molecular markers are applicable for the early screening of low-Cd maize lines and for genetically improving high-Cd maize lines. Of the 13 hybrids, Yuqingzhu23 and Kangnong999 carried Hap_3 and had high grain Cd contents, indicating they may be inappropriate for the production of maize grains intended for human consumption. Molecular marker-assisted breeding may be useful for improving their Cd accumulation characteristics.

## Experimental procedures

### Plant materials and treatments

The association panel comprised 513 maize inbred lines, including 228 lines originating from temperate regions, 258 lines from tropical–subtropical regions, and 27 lines with an admixed origin (Yang et al., 2014). For Cd treatments, maize materials were planted in ZZ and NX. The soil at the study sites was contaminated with Cd because of Cd-rich surface water irrigation. Field trials for the association population were conducted in ZZ (N27°49′50.88″, E113°07′41.23″) and NX (N28°16′51.16″, E112°32′47.13″) in the spring and autumn of 2018. Plants were grown in a single-row plot with two replicates using a randomized complete block design. The F2 population, a BC_1_ segregating population and the parents, *zmcd1-1* and *zmcd1-2* (EMS mutants), as well as the F1 hybrids (obtained by crossing the EMS mutants with test inbred lines) were planted in ZZ in the autumn of 2018. Each row plot had 10 plants in 3 m rows that were separated by 60 cm.

### Soil sampling and composition analysis

Soil samples (0–20 cm depth) were collected from the ZZ and NX study sites using the five-point sampling method (Luo et al., 2017). The soil Cd content was measured by graphite furnace atomic absorption spectrophotometry according to the National Standards of the People’s Republic of China (GB/T 17141-1997). Soil pH was measured using a pH meter.

### Measurement and analysis of maize grain Cd contents

For each maize line replicate, grains were harvested and dried, after which they were ground to a powder. The ground material was passed through a 100-mesh sieve. The collected maize grain powder was weighed and then 0.1–0.5 g was digested with HNO3/HClO4 (9:1, v/v). Finally, the Cd concentration of the digestion solution was determined using the ZEEnit700P atomic absorption spectrometer (Analytikjena, Germany) (Zhao et al., 2018a).

The BLUP data for the GWAS population across environments were estimated via mixed models using the following formula: Y = (1|LINE) + (1|ENV) + (1|REP%in%LINE:ENV) + (1|LINE:ENV) (Y, trait data; “1|”, groups; “:”, interactions; LINE, all testcrosses used; ENV, environments; and REP, replications in one ENV) (Zhao et al., 2018b). The combined ANOVA of four environments was completed and the heritability of the grain Cd content trait was calculated using the ANOVA tool of the IciMapping software (version 4.1) (Luo et al., 2017). The BLUP analysis was performed using the lme4 package of the R software (Zhao et al., 2018b), whereas the correlation analysis was conducted using the Corrplot package of the R software (Luo et al., 2019).

### GWAS analysis and candidate gene prediction

The association panel was genotyped by combining RNA-seq and Illumina MaizeSNP50 BeadChip data (Yang et al., 2014). A total of 1.25 million high-quality SNPs with a minor allele frequency exceeding 5% were used for the GWAS analysis (Yang et al., 2014). The GWAS was completed using TASSEL (version 5) (Bradbury et al., 2007). More specifically, we used a compressed mixed linear (cMLM) model that included the population structure (Q) and the relative kinship (K). A significance threshold of 7.98e-7 (1/independent marker number) was set to control the genome-wide type 1 error rate. The genes within 200-kb flanking regions (± 100 kb of every significant SNP) (Liu et al., 2017) in the B73 RefGen_v2 reference genome in the MaizeGDB database (https://www.maizegdb.org/) were used for predicting candidate genes based on functional annotations.

### BSR-seq and fine mapping

The Jing724 × Mo17 F2 population was used for a BSR-seq analysis. Using the TRIzol reagent method (Luo et al., 2017), total RNA was extracted from the fresh leaves of each F2 seedling grown in a Cd-contaminated field in ZZ. After determining the grain Cd content of the F2 population, the RNA samples of 30 plants with an extremely high grain Cd content and of 30 plants with an extremely low grain Cd content were divided into two equal pools (i.e., high-Cd pool and low-Cd pool). The two RNA pools were sequenced using a HiSeq X instrument (Illumina, USA). The RNA-seq data were deposited in the NCBI Sequence Read Archive database (accession number PRJNA656212) (https://www.ncbi.nlm.nih.gov/bioproject/?term=PRJNA656212.). The RNA-seq reads were trimmed to remove low confidence nucleotides with quality scores < 15 out of 40. The trimmed reads were aligned to the B73 RefGen_v3 reference genome using GSNAP (Wu et al., 2016). The reads uniquely mapped to a single location in the reference genome were used for SNP calling. A Bayesian approach was applied to evaluate the probability of a linkage between each SNP and the causal gene. Associated SNPs were identified based on a probability cutoff of 0.05 (Liu et al., 2012).

For the fine mapping of the major causal locus (*ZmCd1*) identified by BSR-seq, a 5,000-plant BC_1_ population was generated by crossing the Jing724 × Mo17 F1 plants with the Mo17 plants. The causal locus genotype of the high-Cd plants should theoretically be identical to that of Mo17. Thus, we developed new SNP-based KASP markers for the target region with the SNPs identified by BSR-seq. These markers were used to delimit the interval containing *ZmCd1* based on the recombination events in the extremely high-Cd BC_1_ plants (grain Cd content > 0.2 mg/kg). The KASP marker primers were designed and synthesized by LGC Biosearch Technologies (UK) (**Table S3**). The markers were used to determine the genotypes of the BC_1_ plants with the LGC SNPline PCR Genotyping System (UK). Candidate genes were predicted based on functional annotations, gene expression, and target region sequence variations between B73 and Mo17.

### *ZmCd1* sequence analysis

Total DNA was extracted from the leaves of maize inbred line seedlings (e.g., B73, Jing724, D9H, Ky21, CML333, and Mo17) using a published CTAB method (Luo et al., 2019). Primers for amplifying *ZmCd1* (**Table S4**) were designed based on the B73 genome sequence in the MaizeGDB database (https://www.maizegdb.org/). Phanta Max Super-Fidelity DNA Polymerase (Vazyme Biotech Co., Ltd., Nanjing, China) was used for the PCR. The amplified products were sequenced on the ABI3730 sequencer by TsingKe Biological Technology Co. Ltd. (Beijing, China). Sequences were aligned and assembled using the MegAlign software (http://www.freedownload64.com/s/megalign). Transposable elements were predicted for the queried sequences using the CENSOR software (https://www.girinst.org/censor/index.php).

### EMS mutant analysis and allelism test

By screening the Maize EMS induced Mutant Database (http://www.elabcaas.cn/memd/) for the B73 inbred line, we identified *zmcd1-1* (EMS4-038f45) and *zmcd1-2* (EMS4-038f39) as two *ZmCd1* EMS mutants harboring nonsense mutation sites. The mutation sites were validated by sequencing PCR products with specific primers (**Table S4**). The effects of the *zmcd1-1* and *zmcd1-2* mutations on grain Cd accumulation were evaluated by comparing the grain Cd contents of B73 (wild-type), *zmcd1-1*, and *zmcd1-2* plants grown in a Cd-contaminated field in ZZ. The F1 hybrids generated by crossing the high-Cd *zmcd1-1* and *zmcd1-2* mutants with the high-Cd inbred line Mo17 were grown in a Cd-contaminated field in the autumn of 2018. The Cd contents of the kernels produced by the F1 hybrids, Mo17 × *zmcd1-1* and *zmcd1-2* × Mo17, were measured to confirm whether the mutated gene in the *zmcd1-1* and *zmcd1-2* mutants was allelic to the candidate gene for high grain Cd contents in Mo17.

### RT-PCR and qRT-PCR analyses of *ZmCd1*

Total RNA was extracted from whole maize seedlings using the TRIzol reagent (Thermo Fisher Scientific, USA) according to the manufacturer instructions. The extracted RNA was purified with RNase-free DNase 1 (Invitrogen, USA), after which it was used as the template for synthesizing cDNA using the PrimeScript™ II 1st strand cDNA Synthesis Kit (Takara Bio, Beijing, China). The full-length *ZmCd1* cDNA was amplified by PCR using Phanta Max Super-Fidelity DNA Polymerase (Vazyme Biotech). The PCR products were analyzed by 2% agarose gel electrophoresis. The RT-PCR primers are listed in **Table S4**. The PCR products were purified and then sequenced by TsingKe BioTech (Beijing, China). Regarding *ZmCd1^Mo17^*, the RT-PCR products were inserted into the pEASY Blunt Simple vector (TransGen Biotech, Beijing, China) for sequencing.

To analyze *ZmCd1* expression in Jing724 and Mo17, total RNA extracted from seedling roots and leaves at the V3 stage was used to synthesize cDNA as described above. A qRT-PCR assay was performed using the QuantStudio™ 6 Flex system (Thermo Fisher Scientific) and TB Green^^®^^ Premix Ex Taq™ II (Tli RNaseH Plus) (Takara Bio, Beijing, China) with gene-specific primers (**Table S4**). Three biological replicates were used to calculate relative expression levels according to the 2^-ΔΔCt^ method (Luo et al., 2017). The *Actin1* gene (*GRMZM2G126010*) was used as the internal control.

### 5′- and 3′-RACE

The *ZmCd1^Jing724^* and *ZmCd1^Mo17^* transcription start sites were mapped using the FirstChoice™ RLM-RACE Kit (Thermo Fisher Scientific) according to the manufacturer instructions. Briefly, total RNA extracted from Jing724 and Mo17 seedlings was digested with Calf Intestine Alkaline Phosphatase (CIP) to remove uncapped RNA sequences and then treated with Tobacco Acid Pyrophosphatase (TAP) to remove the 5′-cap structure. An RNA adapter was ligated to the CIP/TAP-treated RNA sequences using T4 RNA ligase, after which cDNA was synthesized using M-MLV Reverse Transcriptase. A nested PCR was performed using 5′-RNA adapter primers and gene-specific primers (**Table S4**). A 3′-RACE was performed using total RNA from Ky21 seedlings to map the polyadenylation sites of *ZmCd1^Ky21^* mRNA. First-strand cDNA was synthesized using the 3′-RACE adapter (5′-GCGAGCACAGAATTAATACGACTCACTATAGGT12VN-3′) as a reverse transcription primer. A nested PCR was performed using the 3′-RNA adapter primers and gene-specific primers (**Table S4**). The 5′ - and 3′-RACE PCR products were cloned into the pEASY Blunt Simple vector (TransGen Biotech) and the clones were sequenced by TsingKe Biological Technology Co. Ltd. (Beijing, China).

### Phylogenetic analysis of plant HMAs

The amino acid sequences of 43 previously reported plant HMA proteins were retrieved from the Phytozome database (version 12) (https://phytozome.jgi.doe.gov/pz/portal.html) and the NCBI GenBank database (https://www.ncbi.nlm.nih.gov/genbank/). The sequences included 8 AtHMAs in Arabidopsis (Baxter et al., 2003), 9 OsHMAs in rice (Baxter et al., 2003; Zhiguo et al., 2018), 9 SbHMAs in sorghum (Zhiguo et al., 2018), 12 ZmHMAs in maize (Cao et al., 2019; Zhiguo et al., 2018), BrHMA3 in *B. rapa* (Zhang et al., 2019), HvHMA3-B in barley (Lei et al., 2020), and TdHMA3-B1a in durum wheat (Maccaferri et al., 2019) (**Table S5**). The amino acid sequences of the 43 HMA proteins were aligned using MUSCLE (https://www.ebi.ac.uk/Tools/msa/muscle/). A neighbor-joining phylogenetic tree was constructed using MEGA 7 (Sudhir et al., 2016). Specifically, the Poisson amino acid substitution model and 1,000 bootstrap replicates were used.

### Subcellular localization of ZmCd1

The full-length *ZmCd1^B73^* CDS was fused in frame with the enhanced green fluorescent protein (eGFP)-encoding CDS in the pM999-eGFP vector. The pM999-eGFP empty vector with a 35S-eGFP cassette served as the blank control. A vector containing a sequence encoding the AtTIP2;3-mCherry fusion protein (tonoplast-localized marker) was included in co-transformations. Maize protoplasts were isolated from leaf tissues and co-transformed with the vectors using the plant protoplast preparation and transformation kit (ZhongkeRuitai Biotechnology Co., Ltd., Beijing, China). Fluorescence was detected using the SP8 confocal laser scanning microscope (Leica, USA) with excitation/emission wavelengths of 488 nm/505 nm and 543 nm/610 nm for GFP and mCherry, respectively.

### Development and application of PCR-based functional molecular markers for maize grain Cd accumulation

The genome sequences of 38 maize inbred lines were downloaded from the MaizeGDB database (https://download.maizegdb.org/) to obtain the *ZmCd1* genomic sequences. On the basis of the *ZmCd1* sequence differences among the maize inbred lines, four PCR-based molecular markers (M1–M4) were developed for four regions with varying gene structures (**Figure 5a; Table S6**). Three primers were used in one PCR amplification for the M1–M3 markers to distinguish the InDel variations (**Figure 5a**). For markers M2 and M3, the PCR extension step was less than 1 min to avoid amplifying long transposable elements. Details regarding the primers for these markers are listed in **Table S4**. The PCR products were analyzed by 2% agarose gel electrophoresis. Six haplotypes were detected in the genome-sequenced maize inbred lines based on different M1–M4 marker combinations (**Table S6**). Additionally, 36 elite inbred lines and 13 hybrid varieties (**Table S7**) commonly cultivated in China were grown in a Cd-contaminated field in ZZ in the autumn of 2018. The CTAB method (Luo et al., 2019) was used to extract the DNA from these materials. The M1–M4 markers were used for genotyping the inbred lines and hybrid varieties to predict the grain Cd contents.

## Acknowledgments

We thank Prof. Jianbing Yan for kindly providing the GWAS panel. This research was financially supported by the Beijing Municipal Natural Science Foundation (6204041), the Youth Research Fund of Beijing Academy of Agriculture and Forestry Sciences (QNJJ202028), the National Key Research and Development Program of China (2016YFD0300109), the XiangCaiNongZhi (2019-33), the Construction of Collaborative Innovation Center of Beijing Academy of Agricultural and Forestry Sciences (Collaborative Innovation Center of Crop Phenomics, KJCX201917), and the Beijing Scholars Program (BSP041).

## Authors’ contributions

Z.Y. (Yanxin Zhao), T.B., C.Z. (Zhihui Chen), and Z.J. (Jiuran Zhao) conceived the study. Z.Y. (Yanxin Zhao), T.B., L.M., and S.W. designed the experiments. T.B., L.M., Z.Y. (Yunxia Zhang), G.H., L.J., Z.R., F.Z., K.M., and L.H. performed the experiments. Z.Y. (Yanxin Zhao), L.D., and Z.J. (Jianhua Zhang) analyzed the BSR-seq data. L.M., Z.Y. (Yanxin Zhao), C.Z. (Zhongyang Cao), W.R., and W.Y. analyzed the experimental data. L.M., and Z.Y. (Yanxin Zhao) wrote the manuscript. All authors revised and approved the final manuscript, and agreed to be accountable for this study.

## Supporting information

**Figure S1. Frequency distribution of maize grain Cd contents in GWAS plants in four environments.** ZZC, spring in ZZ. ZZQ, autumn in ZZ. NXC, spring in NX. NXQ, autumn in NX.

**Figure S2. GWAS of maize grain Cd accumulation in four environments.** (a–f) Manhattan plots for grain Cd accumulation in four environments and the best linear unbiased prediction (BLUP) data. The red horizontal line represents the significance cutoff (*P* = 7.97e-7). NX, Ningxiang. (g–l) Quantile-Quantile plots for the GWAS MLM + Q + K model. BLUP4EN, BLUP values for the grain Cd contents in four environments. BLUPZZ, BLUP values for the grain Cd contents in two seasons in Zhuzhou (ZZ). NXC, spring in Ningxiang (NX). NXQ, autumn in NX. ZZC, spring in ZZ. ZZQ, autumn in ZZ.

**Figure S3. *ZmHMA3* and *ZmHMA4* expression levels in diverse tissues.** Relative *ZmHMA3* (a) and *ZmHMA4* (b) expression levels in diverse tissues based on the data in the Maize eFP browser (http://bar.utoronto.ca/efp_maize/cgi-bin/efpWeb.cgi). Color intensity from yellow to red represents the expression levels (low to high, respectively). (c) Differential expression analysis of *ZmHMA3* and *ZmHMA4*. The linearized RMA expression signals in 60 diverse tissues were downloaded from the Maize eFP browser (http://bar.utoronto.ca/efp_maize/cgi-bin/efpWeb.cgi). Error bar = standard deviation.

**Figure S4. Expression of *ZmHMA4* gene in B73, Jing724 and Mo17.** RT-PCR product electrophoresis of *ZmHMA4* were run on 2% agarose gel. Root and leaf RNAs of B73, Jing724 and Mo17 were isolated for cDNA synthesis. RT-PCR was performed with gene-specific primers HMA61F and HMA2564R shown in **Table S4**. Maize *Actin1* gene was amplified as an internal control with the primers actinF2 and actinR2. M, DL2000 DNA Marker (Beijing TsingKe Biotech Co., Ltd, China). -RT, cDNA synthesized without reverse transcriptase. The gene-specific PCR products were indicated by the orange arrows.

**Figure S5. Variations in the *ZmCd1^Mo17^* transcript.** (a) Number of RNA-seq reads mapped to the *ZmCd1* gene in Mo17. The RNA-seq reads of the high-Cd and low-Cd pools were mapped to the Mo17 reference genome. The read counts for *ZmCd1^Mo17^* were visualized using the IGV browser (https://software.broadinstitute.org/software/igv/). The *ZmCd1^Mo17^* gene structure was generated based on the *ZmCd1^B73^* coding sequence. (b) Structures of all 19 *ZmCd1^Mo17^* transcripts. Transcript sequences were determined by sequencing 109 cDNA clones, which were constructed from RT-PCR products using the 7F and 7R primers and Mo17 leaf cDNA as the template. The number of clones and the length of the PCR products for each transcript are indicated in parentheses. The transposable element-inserted regions in introns 1 and 3 are highlighted in red.

**Figure S6. Alignment of HMA3 proteins from maize, sorghum, Arabidopsis, rice, wheat, and barley.** The amino acid sequences of the following proteins were aligned using Clustal Omega (https://www.ebi.ac.uk/Tools/msa/clustalo/) and displayed using GeneDoc (http://nrbsc.org/gfx/genedoc): ZmHMA3 (GRMZM2G175576), SbHMA3a (Sobic.002G083000), AtHMA3 (AT4G30120), OsHMA3 (LOC_Os07g12900), TdHMA3-B1a (NCBI GenBank ID: KF683293), and HvHMA3-B (NCBI GenBank ID: LC523825). The conserved domains were predicted using the Pfam database (http://pfam.xfam.org/) and are framed by green, red, and blue lines. Green box, heavy metal-associated domain (PF00403). Red box, E1–E2 ATPase domain (PF00122). Blue box, haloacid dehalogenase-like hydrolase (PF00702).

**Figure S7. Grain, stem, and leaf Cd contents of five maize inbred lines and six maize hybrid varieties.** The inbred lines and hybrids were grown in a Cd-contaminated field in Zhuzhou in the autumn of 2018. After harvesting, the grains, stems, and leaves were collected and dried before measuring their Cd contents. n = 3, error bar = standard error.

**Table S1. Summary of the QTL related to the maize grain Cd content identified in a GWAS.**

**Table S2. List of the SNPs on chromosome 2 significantly associated with the grain Cd content.**

**Table S3. SNP-based KASP markers used for fine mapping *ZmCd1.***

**Table S4. List of the PCR primers used in this study.**

**Table S5. List of plant HMA-encoding genes used for constructing a phylogenetic tree.**

**Table S6. Information regarding the PCR-based markers in maize inbred lines with sequenced genomes.**

**Table S7. Maize inbred lines and hybrid varieties used in this study.**

**Supplemental Dataset S1. Alignment of ZmHMA3 from B73, D9H, CML333, Mo17, and Ky21.aln**

