## Supplemental Figure S1-S7 for "Natural variations in the P-type ATPase heavy metal transporter ZmCd1 controlling cadmium accumulation in maize grains"

**Supplementary Figures**

**Figure S1. Frequency distribution of maize grain Cd contents in GWAS plants in four environments.**

**Figure S2. GWAS of maize grain Cd accumulation in four environments.**

**Figure S3. *ZmHMA3* and *ZmHMA4* expression levels in diverse tissues.**

**Figure S4. Expression of *ZmHMA4* gene in B73, Jing724 and Mo17.**

**Figure S5. Variations in the $ZmCd1^{Mo17}$ transcript.**

**Figure S6. Alignment of HMA3 proteins from maize, sorghum, Arabidopsis, rice, wheat, and barley.**

**Figure S7. Grain, stem, and leaf Cd contents of five maize inbred lines and six maize hybrid varieties.**

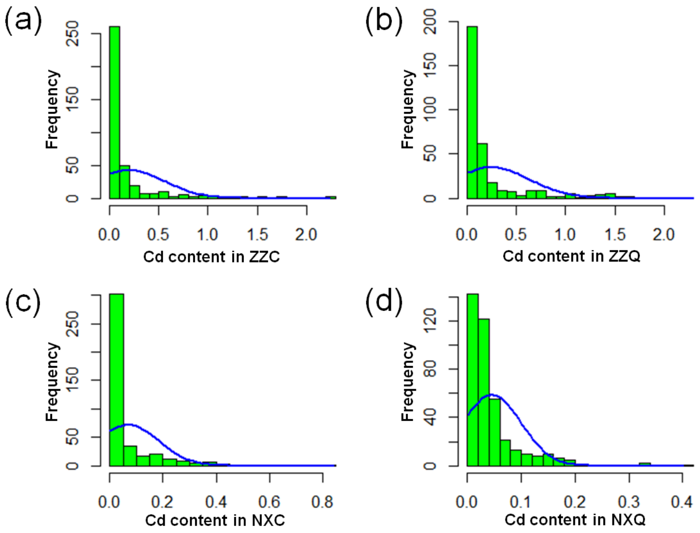


**Figure S1. Frequency distribution of maize grain Cd contents in GWAS plants in four environments.** ZZC, spring in ZZ. ZZQ, autumn in ZZ. NXC, spring in NX. NXQ, autumn in NX.

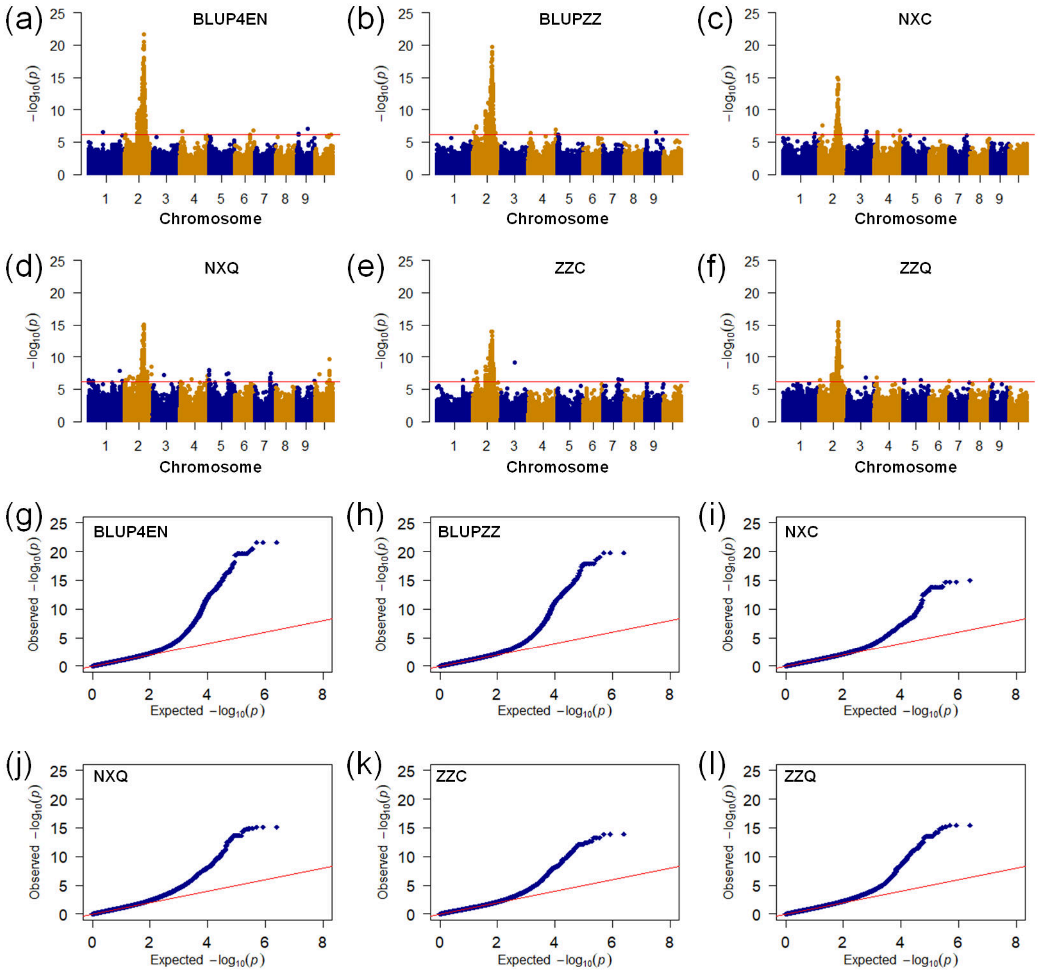


**Figure S2. GWAS of maize grain Cd accumulation in four environments.** (a–f) Manhattan plots for grain Cd accumulation in four environments and the best linear unbiased prediction (BLUP) data. The red horizontal line represents the significance cutoff ($P$ = 7.97e-7). NX, Ningxiang. (g–l) Quantile-Quantile plots for the GWAS MLM + Q + K model. BLUP4EN, BLUP values for the grain Cd contents in four environments. BLUPZZ, BLUP values for the grain Cd contents in two seasons in Zhuzhou (ZZ). NXC, spring in Ningxiang (NX). NXQ, autumn in NX. ZZC, spring in ZZ. ZZQ, autumn in ZZ.

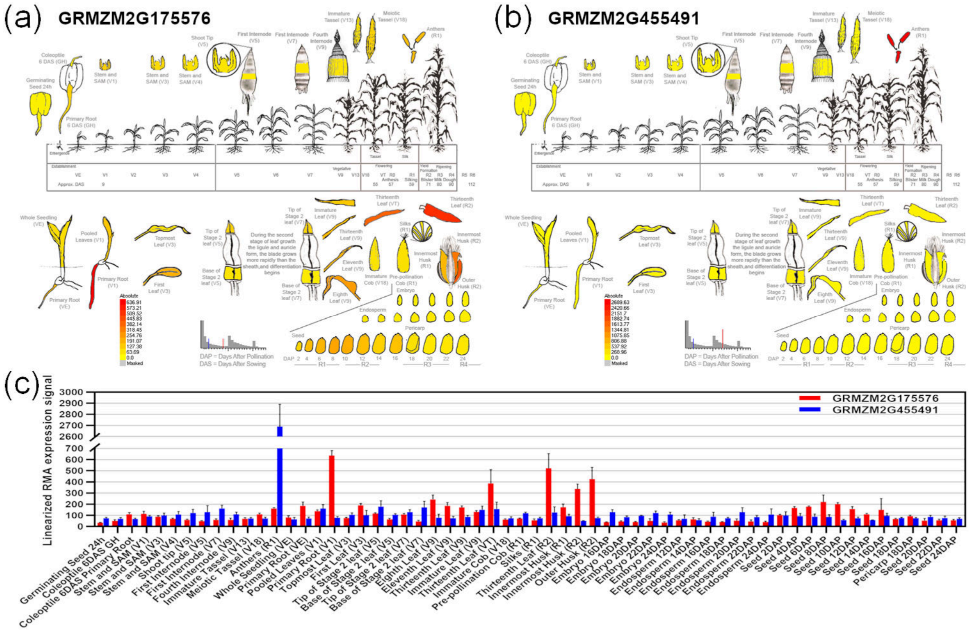


**Figure S3. *ZmHMA3* and *ZmHMA4* expression levels in diverse tissues.** Relative *ZmHMA3* (a) and *ZmHMA4* (b) expression levels in diverse tissues based on the data in the Maize eFP browser (http://bar.utoronto.ca/efp_maize/cgi-bin/efpWeb.cgi). Color intensity from yellow to red represents the expression levels (low to high, respectively). (c) Differential expression analysis of *ZmHMA3* and *ZmHMA4*. The linearized RMA expression signals in 60 diverse tissues were downloaded from the Maize eFP browser (http://bar.utoronto.ca/efp_maize/cgi-bin/efpWeb.cgi). Error bar = standard deviation.

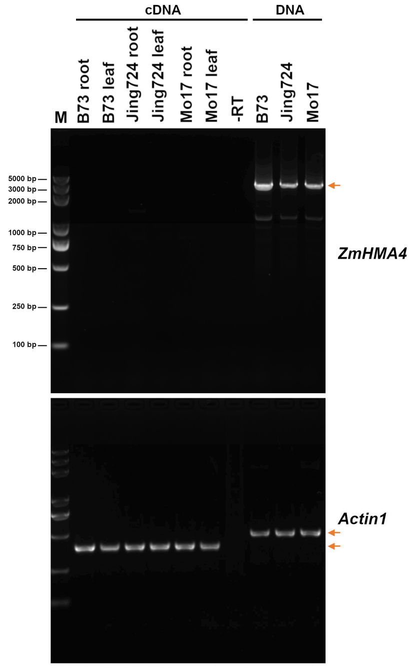


**Figure S4. Expression of *ZmHMA4* gene in B73, Jing724 and Mo17.** RT-PCR product electrophoresis of *ZmHMA4* were run on 2% agarose gel. Root and leaf RNAs of B73, Jing724 and Mo17 were isolated for cDNA synthesis. RT-PCR was performed with gene-specific primers HMA61F and HMA2564R shown in **Table S4**. Maize *Actin1* gene was amplified as an internal control with the primers actinF2 and actinR2. M, DL2000 DNA Marker (Beijing TsingKe Biotech Co., Ltd, China). -RT, cDNA synthesized without reverse transcriptase. The gene-specific PCR products were indicated by the orange arrows.

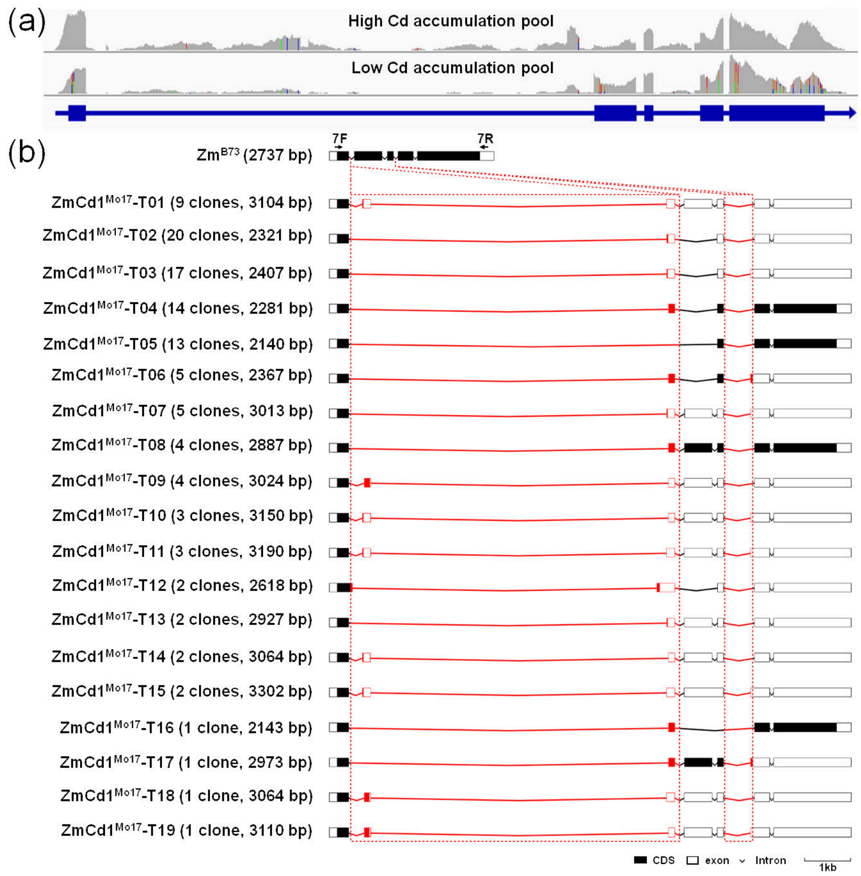


**Figure S5. Variations in the *ZmCd1$^{Mo17}$* transcript.** (a) Number of RNA-seq reads mapped to the *ZmCd1* gene in Mo17. The RNA-seq reads of the high-Cd and low-Cd pools were mapped to the Mo17 reference genome. The read counts for *ZmCd1$^{Mo17}$* were visualized using the IGV browser (https://software.broadinstitute.org/software/igv/). The *ZmCd1$^{Mo17}$* gene structure was generated based on the *ZmCd1$^{B73}$* coding sequence. (b) Structures of all 19 *ZmCd1$^{Mo17}$* transcripts. Transcript sequences were determined by sequencing 109 cDNA clones, which were constructed from RT-PCR products using the 7F and 7R primers and Mo17 leaf cDNA as the template. The number of clones and the length of the PCR products for each transcript are indicated in parentheses. The transposable element-inserted regions in introns 1 and 3 are highlighted in red.

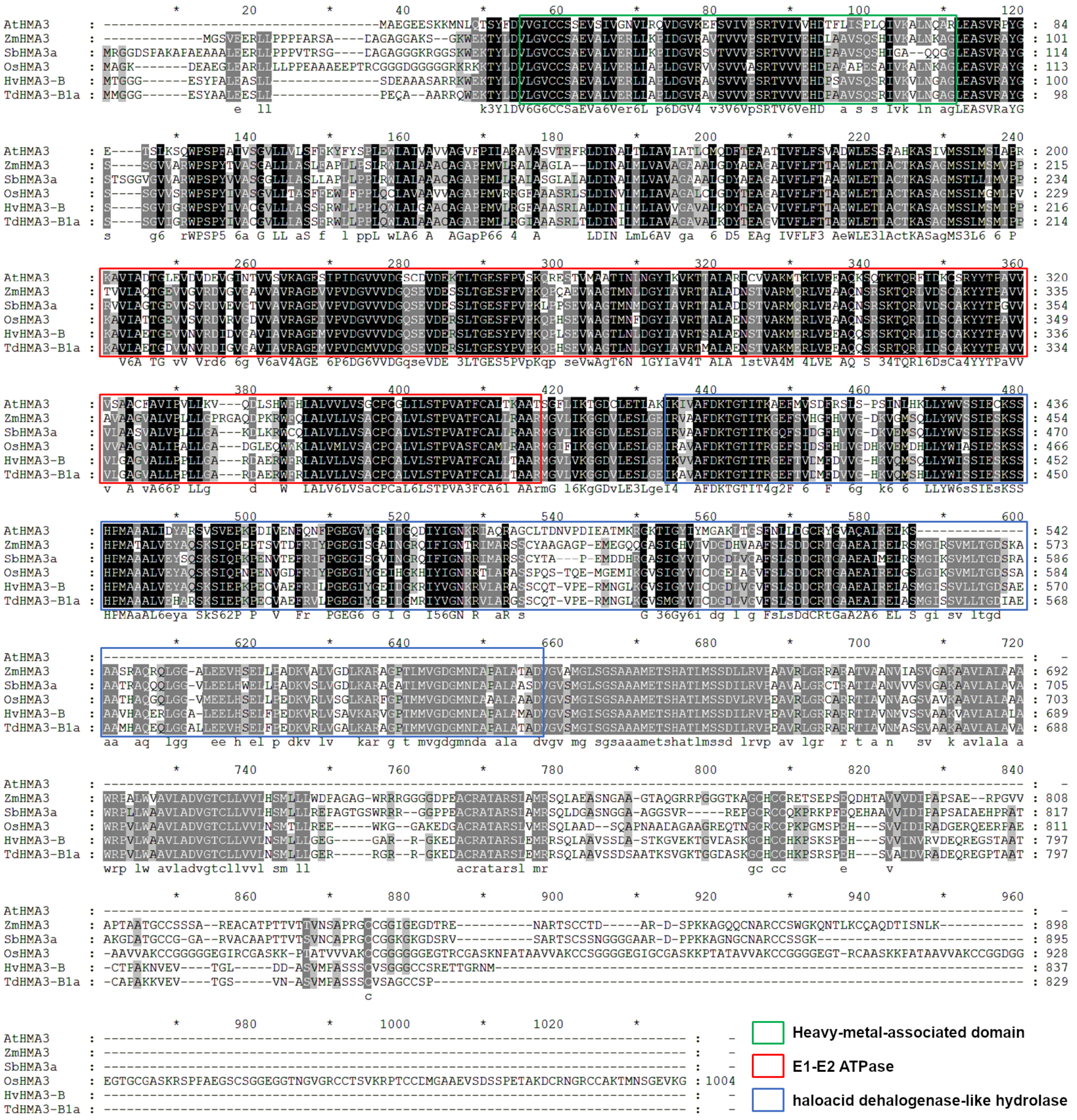


**Figure S6. Alignment of HMA3 proteins from maize, sorghum, Arabidopsis, rice, wheat, and barley.** The amino acid sequences of the following proteins were aligned using Clustal Omega (https://www.ebi.ac.uk/Tools/msa/clustalo/) and displayed using GeneDoc (http://nrbsc.org/gfx/genedoc): ZmHMA3 (GRMZM2G175576), SbHMA3a (Sobic.002G083000), AtHMA3 (AT4G30120), OsHMA3 (LOC_Os07g12900), TdHMA3-B1a (NCBI GenBank ID: KF683293), and HvHMA3-B (NCBI GenBank ID: LC523825). The conserved domains were predicted using the Pfam database (http://pfam.xfam.org/) and are framed by green, red, and blue lines. Green box, heavy metal-associated domain (PF00403). Red box, E1–E2 ATPase domain (PF00122). Blue box, haloacid dehalogenase-like hydrolase (PF00702).

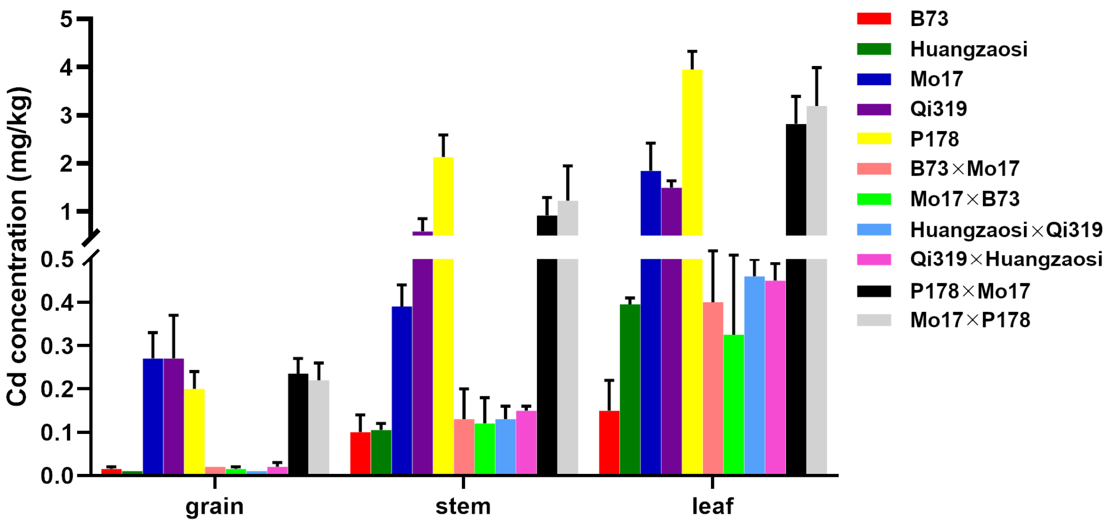


**Figure S7. Grain, stem, and leaf Cd contents of five maize inbred lines and six maize hybrid varieties.** The inbred lines and hybrids were grown in a Cd-contaminated field in Zhuzhou in the autumn of 2018. After harvesting, the grains, stems, and leaves were collected and dried before measuring their Cd contents. n = 3, error bar = standard error.
